# *Schistosoma mansoni*-induced host immunopathology is gut microbiota dependent

**DOI:** 10.1101/2025.01.24.634684

**Authors:** Anna Overgaard Kildemoes, Gabriele Schramm, Bente Pakkenberg, Fynn Krause, Morten Arendt Rasmussen, Camilla Hartmann Friis Hansen, Witold P. Kot, Anne Majgaard Jensen, Jakub Wawrcyniak, Julian L. Griffin, Shona Wilson, Søren Skov, Axel Kornerup Hansen, Dennis Sandris Nielsen, Birgitte Jyding Vennervald

## Abstract

Despite progress in schistosomiasis control during recent decades in endemic areas, this parasitic blood-fluke infection continues to pose a substantial public health burden. In depth understanding of the mechanisms behind the complex immunopathology and drivers of morbidity in schistosomiasis is urgently needed. Chronic infection with the parasitic blood fluke *Schistosoma mansoni* manifests the most severe pathology in relation to egg-induced host immune responses and fibrosis development. Host immune homeostasis is influenced by the commensal gut microbiota, which may therefore also affect systemic immunopathology induced by *S. mansoni.* Both humans and experimental animal models show metabolomic changes related to gut microbial and liver metabolism during *S. mansoni* infection, further supporting a link between gut microbiota and regulation of *S. mansoni* pathology. To investigate whether a radically changed gut microbiota composition would result in changed *S. mansoni* egg-induced pathology, a mouse model combining antibiotic treatment and infection was established. In the model, the commensal gut microbial composition of female C57BL/6-NTac mice was altered by oral administration of a broad-spectrum ampicillin-vancomycin cocktail prior to infection with *S. mansoni* and throughout the experiment. An ecosystem view of the host immune milieu was collated by measuring a wide range of parameters: gut microbiome characterisation, immune profiling on the transcriptional level (ileum and liver) supported by flow cytometry of cells (spleen and mesenteric lymph nodes) and bead-based measurements of immune parameters in sera, short-chain fatty acid measurements in sera and caecal content, intestinal permeability assessment, liver enzyme levels in sera, and histology and stereology to characterise immunopathology (ileum and liver). The application of stereological principles to address the non-spherical nature of egg-induced liver granulomas and ileum inflammation supported by liver enzyme levels, collagen deposition, and immune parameters, enabled demonstration of significantly less granuloma formation in livers from antibiotics treated mice compared to controls. The observed difference in the degree of egg-induced inflammatory response in livers from infected mice mediated by antibiotics treatment, was not observed in ileum tissue and could not be explained by infection burden. Our mouse model results demonstrate a role for the gut microbiota and intestinal immune milieu in regulation of systemic *S. mansoni* infection-driven liver pathology and, hence, potential morbidity. In translational context, these findings warrant that drivers of gut microbial changes and intestinal immune milieu must be considered when pursuing novel holistic interventions for treatment or alleviation of persistent schistosomiasis.

## Introduction

Infection with the parasitic blood fluke *Schistosoma mansoni* remains a substantial poverty-associated public health challenge in endemic regions (1). Most of these regions are located in Sub-Saharan Africa, where more than 50 million people are infected (2–4). Despite large-scale control programmes, focal infection hotspots with persistently infected people and hence high morbidity prevail (5).

The pathology caused by *S. mansoni* is essentially host immune response driven and comprises both host protective and damaging elements. Pairs of adult schistosomes lay eggs directly in the mesenteric veins, after which the eggs traverse the intestinal barrier to leave the host with faecal matter for subsequent propagation of transmission (6). The eggs secrete an array of pro-inflammatory and immunomodulatory factors which initiate transport of eggs across host tissue barriers (7). A large proportion of eggs do not successfully cross the intestinal barrier but are instead swept with the bloodstream and lodge in other tissues particularly in the liver, where they induce eosinophil rich granulomas (8). These granulomas encapsulate the schistosome eggs and play a dual role, where they both protect the host from hepatotoxic substances secreted by the *S. mansoni* eggs (9, 10), as well as cause severe disease. Severe disease occurs particularly in chronic high intensity infections typically by development of liver fibrosis resulting in portal hypertension and oesophageal varices (11, 12). Host immune homeostasis is relevant for understanding complex manifestations of differences in degree of morbidity and underlying pathology induced by infection with *S. mansoni.* Recently, much focus has been on the role of the commensal gut microbial composition and short-chain fatty acid (SCFA) availability on host immune homeostasis in many disease models and clinical contexts (13–17), which points to a potential link to immunopathology induced by *S. mansoni*. Interestingly, in mice it has been demonstrated that depletion of gut microbiota in *S. mansoni*-infected animals resulted in a reduction of intestinal inflammation and reduced granuloma formation in response to eggs lodged in intestinal tissue (18). However, the effect was local and no differences were observed in liver tissue. Both humans and mice show changes in molecular markers associated with liver metabolism and gut microbial activity in response to hepato-intestinal schistosome infection observed by metabolomic approaches (19–21). These studies implicate a role for gut microbiota-mediated immune regulation in *S. mansoni*-induced immunopathology.

A range of factors such as diet, infections, drugs, and other environmental factors can influence gut microbial composition in humans and experimental animal models (22–25). Here we hypothesised, that a strong alteration of the gut microbial composition *prior* to a primary infection with *S. mansoni* can affect local and systemic host immune responses, including development of immunopathology such as liver granulomas. The study takes a host-parasite-gut microbiota ecosystem view and describes host immune response profiles, faecal microbiome composition, SCFA levels and degree of induced immunopathology in an infection mouse model combining antibiotics-mediated alteration of gut microbial composition and *S. mansoni* infection. Our findings underline the necessity to take gut microbial composition and hence gut immune homeostasis into account when evaluating immune responses including immunopathology in *S. mansoni* models as well as in human disease contexts. This is essential to understand drivers behind persistent and hard-to-treat immunopathology and complex morbidity of *S. mansoni* infections.

## Results

### *S. mansoni* infection drive a change in gut microbiome composition

A *S. mansoni* infection model incorporating alteration of the gut microbial composition mediated by a broad-spectrum ampicillin and vancomycin cocktail was established. It consisted of four groups (n = 10 each) of adult seven weeks old female C57BL/6-NTac mice; control (Ctr), antibiotics-treated (Abx), *S. mansoni*-infected (Sm) and antibiotics-treated + *S. mansoni*-infected (AbxSm). The antibiotics were administered to mice prior to infection and throughout the experiment. To confirm that *S. mansoni* infection would result in changes in host gut microbial composition and determine efficacy of the antibiotics cocktail, faecal samples were collected longitudinally from individual mice. Samples were obtained at baseline, at a prepatent time point before parasite egg production onset, and at a patent time point when the parasites lay eggs that penetrate the gut wall as well as lodge predominantly in the liver (Figure 1 and 2). We observed a significantly higher gut microbial species richness (α-diversity) for Ctr mice compared to Sm mice at the patent time point (Suppl. Figure S2B), whereas the Shannon effective index indicated no differences between the two groups (Figure 1A). In contrast, the α-diversity was as expected markedly lowered after antibiotics treatment (Figure 2A), but infected (AbxSm) mice showed a higher number of different species compared to non-infected (Abx) mice at both prepatent and patent time points (Suppl. Figure S1D). Infection drives the faecal microbiome compositional changes both at the prepatent and patent time point as seen on Figure 1B. Similarly, gut microbial compositional (β-diversity) differences between infected and non-infected mice were also noticeable during antibiotics treatment at the patent time point (Figure 2B). The observed differences in faecal microbiome composition were small, albeit representing changes in genera of known physiological relevance (Suppl. Figure S1A and S2AB). Infection was associated with elevated relative abundance of *Bacteroides* spp., while the *Rikenellaceae* RC9 group, *Alistipes* spp. and *Eubacterium* spp. were lowered or not detected in infected mice. Various clostridia were also associated with infection, particularly at the patent time point where *Marvinbryntia* spp. and the *Clostridiales vadin* BB60 group relative abundance were elevated compared to controls. Additionally, the relative abundance of *Prevotella* was lower at the prepatent time point for infected mice compared to controls, but increased at the patent time point. The opposite pattern was observed for *Lactobacillus* sp. Schistosome infection also promoted higher levels of *Clostridiales vadin* BB60 group in the antibiotics-treated mice compared to their controls (Figure 2C, Suppl. Figure S2B) and one of the cages of AbxSm mice presented with elevated *Verrumicrobiaceae* and *Fusobacteriaceae* members at the patent time point.

**Figure 1:**
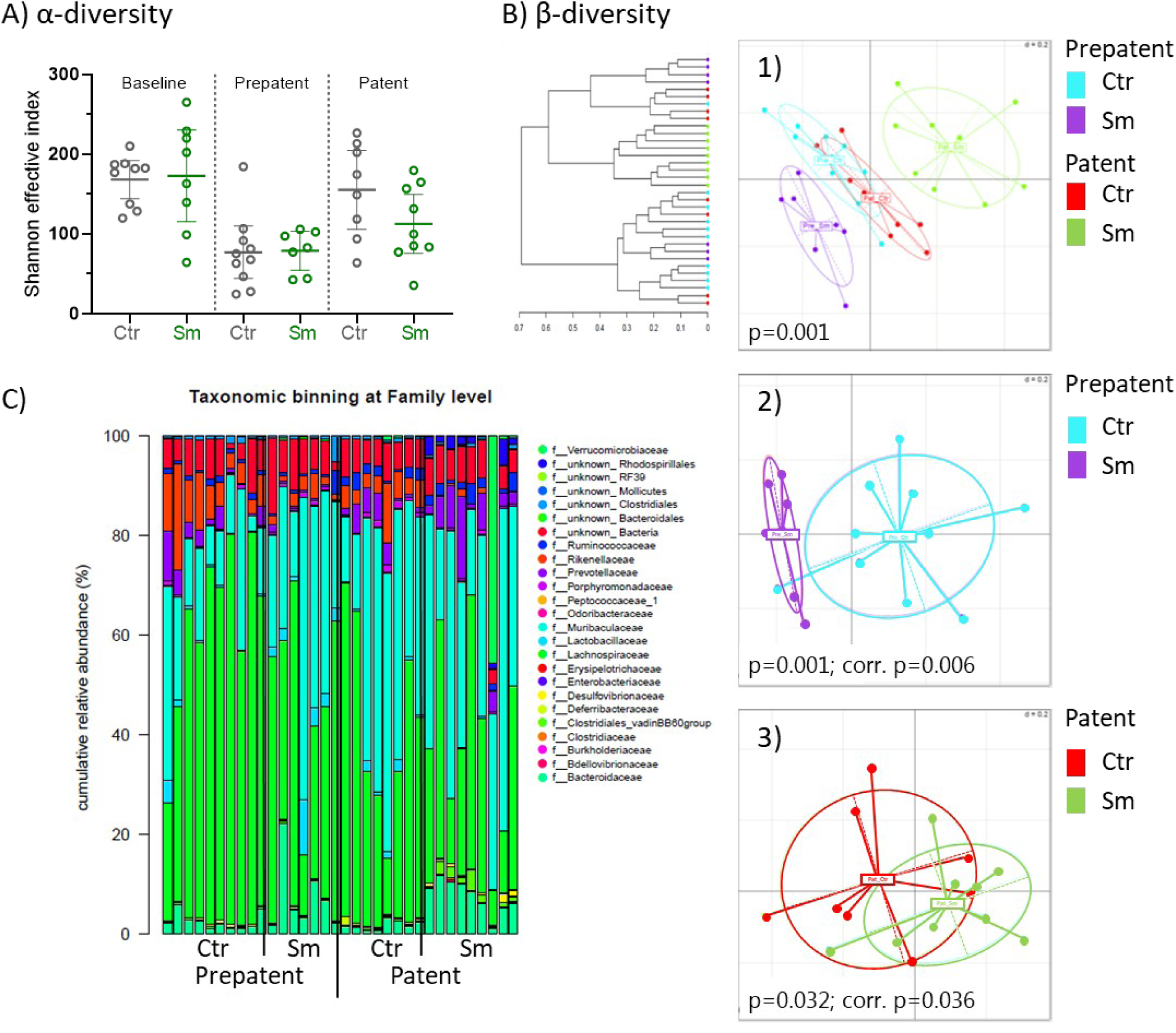
Patent *S. mansoni* infection in mice drive gut microbiome composition change. Faecal microbiome characterisation based on 16S rRNA gene amplicon sequencing of control (Ctr) and *S. mansoni*-infected (Sm, 120 cercariae) mice at 26 days (prepatent: Ctr n=10, Sm n=7. Mice were 12 weeks old) and 41 days (patent: Ctr n=8, Sm n=9. Mice were 14 weeks old) post-infection time points. A) Microbial diversity (Shannon effective α-diversity index; species richness can be found in supplementary figure 1B). B) Microbial compositional differences (β-diversity) shown as meta-NMDS plot 1) of both groups at prepatent and patent time points and the corresponding hierarchical clustering dendrogram based on Ward’s minimum variance method and NMDS plots of pairwise comparisons at 2) prepatent and 3) patent time point. NMDS plots were generated based on generalised UniFrac distance metrics and p values shown are determined by PERMANOVA analysis. C) Gut microbiome composition at taxonomic family level. *Schistosoma* spp. specific qPCR was used to confirm that time points were respectively *Schistosoma* spp. DNA negative and positive for the Sm group. One group of Sm mouse was found dead day 26 post-infection and excluded from analysis. Additional relevant differences in relative abundance between groups are shown in Suppl. Figure S1A and S2A.

**Figure 2:**
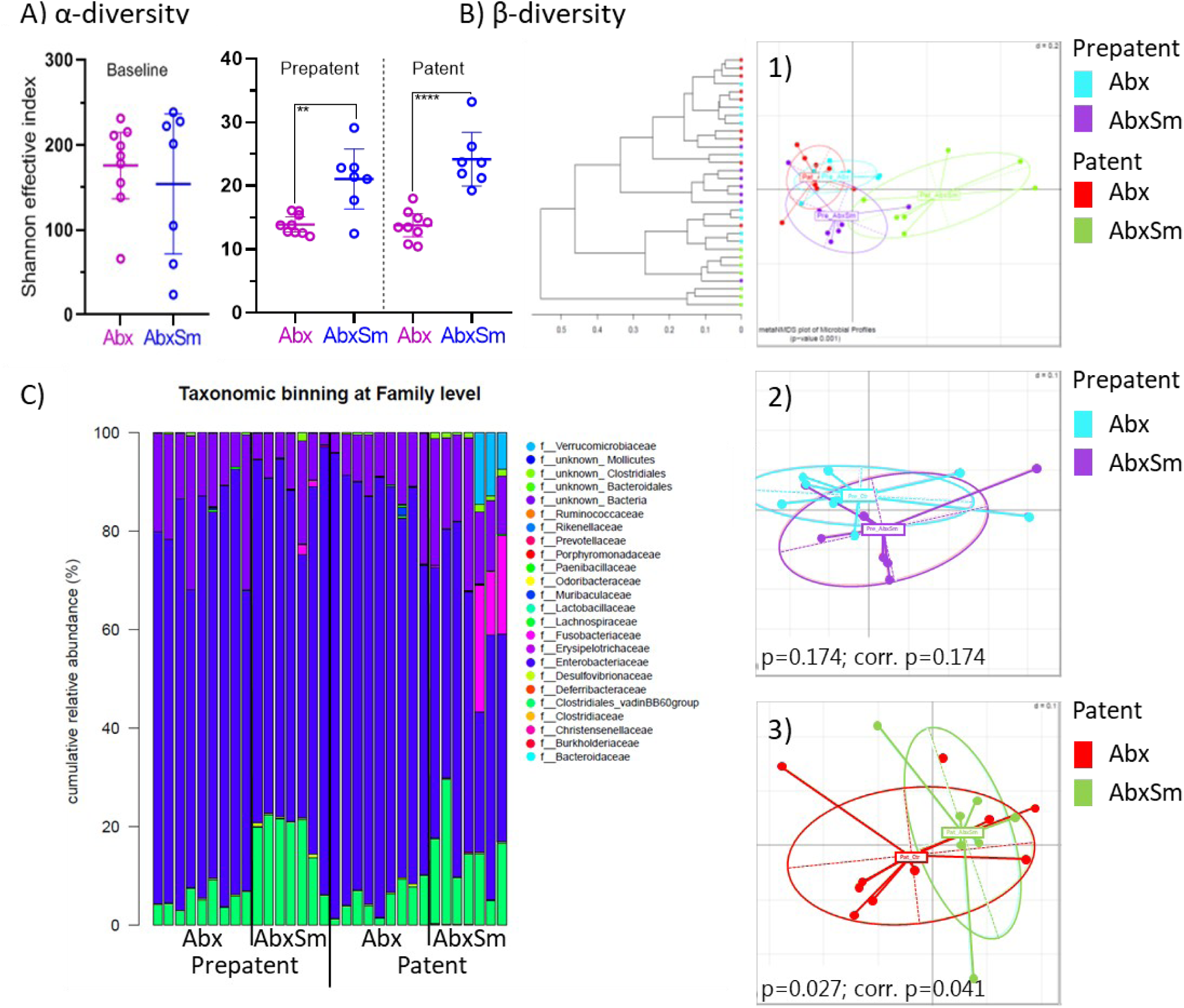
*S. mansoni* infection drive a change in gut microbiome composition detectable in faeces also in antibiotics-treated mice. Faecal microbiome characterisation based on 16S rRNA gene amplicon sequencing of antibiotics-treated (Abx) and antibiotics-treated *S. mansoni*-infected (AbxSm, 120 cercariae) mice at 26 days (prepatent: Abx n=9, AbxSm n=7) and 41 days (patent: Abx n=9, AbxSm n=7) post-infection time points. A) Microbial diversity (Shannon effective α-diversity index, species richness can be found in Supplementary Figure 1CD). B) Microbial compositional differences (β-diversity) shown as meta-NMDS plot 1) of both groups at prepatent and patent time points and the corresponding hierarchical clustering dendrogram based on Ward’s minimum variance method and NMDS plots of pairwise comparisons at 2) prepatent and 3) patent time point. NMDS plots were generated based on generalised UniFrac distance metrics and p values shown are from PERMANOVA analysis. C) Gut microbiome composition at taxonomic family level. *Schistosoma* spp. specific qPCR was used to confirm that time points were respectively *Schistosoma* spp. DNA negative and positive for the Sm group. Three group AbxSm mice were found dead day 33 post-infection and one group Sm at day 26 post infection. Additional relevant differences in relative abundance between groups are shown in suppl. Figure 1A and 2B.

### Alteration of gut microbial composition affects degree of egg-induced liver pathology

Since the gut microbiota composition is important for many immunological processes, we next questioned whether immunopathology elicited by *S. mansoni* eggs would be impacted by the antibiotics-mediated microbiome alteration. Traditional methods evaluating the sizes of *S. mansoni* egg-induced granulomas in tissues are based on two dimensional measurements (26). However, schistosomiasis-associated granulomas are not of a spherical nature, so in order to obtain more precise estimates of the degree of egg-induced inflammation in liver and ileum tissues, we used stereology (Suppl. Figure S3). The advantages of this approach are the incorporation of random sampling and a three-dimensional aspect for unbiased quantification (27, 28). Hepatomegaly in *S. mansoni*-infected compared to uninfected mice was observed based on simple organ weight (Suppl. Figure S4) corresponding well to volumes determined by stereology for entire livers (Figure 3A, p<0.001). Furthermore, the estimated proportion of liver volume consisting of granulomatous tissue was significantly lower in AbxSm mice compared to mice solely infected (p=0.007, Figure 3C). In contrast, despite a clear effect of infection on ileum volume (p<0.01), no differences were observed between the Sm and AbxSm groups neither in terms of ileum enlargement nor proportion of inflamed tissue (Figure 3E, G). The *S. mansoni* infection burden was similar in the two infected groups both in terms of numbers of worms and eggs and cannot explain the observed effect of altering the gut microbial composition on the degree of granulomatous inflammation in the liver (Suppl. Figure S5). Linear regression fits revealed no differences in terms of the positive relationship between number of eggs present and volume or proportion of inflammation in the ileum (Figure 3F, H) between the two infected groups.

**Figure 3:**
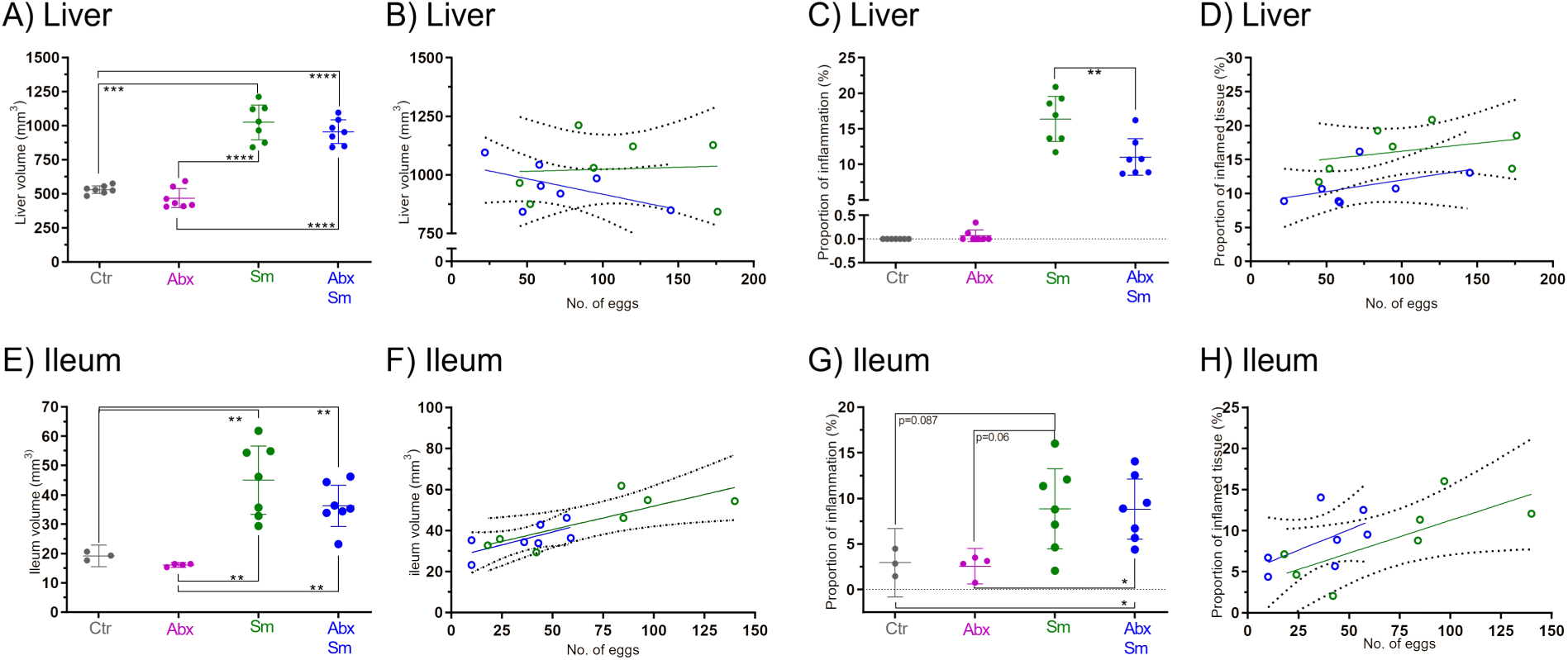
Volume and relative degree of *S. mansoni-*induced inflammation determined by stereological principles in liver and ileum tissues. The estimated A) entire liver volumes (mm^3^) and C) proportion of granulomatous tissue in liver (%) per animal (n=7 in all groups). For ilea the estimated volume (mm^3^) is given in E) based on 36 cross sections per animal and the G) proportion of inflamed tissue in ileum (%) for control (Ctr, n=3), antibiotics-treated (Abx, n=4 ileum), *S. mansoni*-infected (Sm, n=7) and antibiotics-treated and *S. mansoni*-infected (AbxSm, n=7). C) The clearly significant differences between uninfected and infected groups are not indicated on graph. Linear regression plots showing the association between volume of tissue (B, F) and the degree of inflammation (D, H) in relation to the total number of eggs observed in counting frames for all liver sections per animal (B, D) and the total number of eggs observed in 36 ileum cross sections per animal (F, H). Only the elevation on D) is significantly different between Sm and AbxSm (p=0.026). Infected groups were exposed to 120 cercariae per mouse. Bars indicate means with 95% CI, Welch’s ANOVA, Dunnett’s multiple correction.

However, no relation between number of eggs and the proportion of granulomatous tissue was found in the livers of infected mice (Figure 3B, D), underlining the observed difference in local relative degree of inflammation. As infection intensity was comparable, we questioned the activity of the eggs lodged in tissues and therefore assessed whether the eggs had similar excretory/secretory capacity in the two infected groups. Both in liver and ileum a mix of mature and immature eggs were examined by staining for the schistosome egg antigens IPSE/alpha-1 and omega-1, which are secreted by mature eggs (Figure 4C). No differences in the presence of antigens associated with eggs or the diffusion of the antigen in the granulomas surrounding the eggs were seen between the Sm and AbxSm groups. The different staining intensity of IPSE/alpha-1 and omega-1 reflects the presence of these antigens in the egg secretions with IPSE/alpha-1 representing more than 80% of the secretory products (9, 73, 74) (Figure 4C). In contrast to the egg antigens, different numbers of collagenous depositions associated with granulomas and small vessels were seen in the liver (based on a single slide/liver, Figure 4AB). Figure 4D shows representative pictures of eggs travelling across the intestinal wall in a clustered row. No differences in terms of pathology were observed in relation to egg migration between the Sm and AbxSm groups. Multiple examples of both clustered and single eggs were observed without marked granuloma formation in ilea.

**Figure 4:**
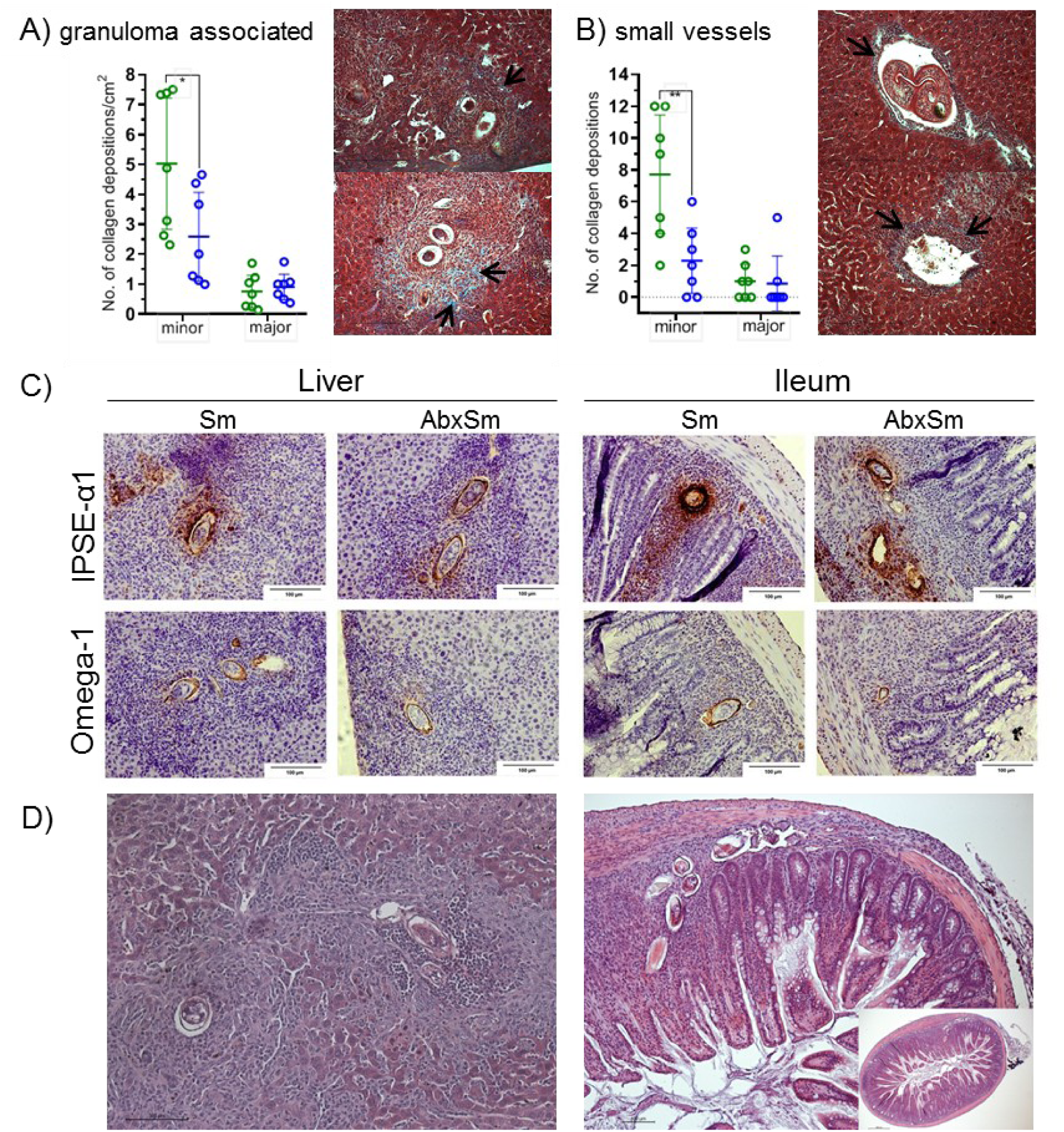
Collagen deposition and egg antigen secretion in liver and ileum tissues. The number of collagen depositions categorised and counted in a blinded manner as “minor” or “major” depositions is shown for A) granuloma-associated (p=0.043) and B) small vessels-associated (p=0.009) depositions based on a single large liver section/animal (unpaired, two-tailed t-tests). *S. mansoni*-infected (Sm) are shown to the left in green and antibiotics-treated and *S. mansoni*-infected (AbxSm) to the right in blue on graphs. Representative pictures of collagen depositions (examples shown by black arrows) in A) granuloma-associated and B) small vessel-associated liver sections shown by Masson’s trichrome staining (20x, scale bar 200 µm) where the top pictures are “minor” and bottom pictures are “major” depositions. C) Representative pictures of localisation and excretion of the egg antigens IPSE/alpha-1 and omega-1 from eggs in liver and ileum sections (scale bars are 100 µm). Intensity of stain relates to antigen quantity. No differences between Sm and AbxSm groups were observed qualitatively. D) Examples of liver granulomas and clustered egg migration through ileum tissue (10x and 4x objectives, scale bars of 100 µm and 400 µm respectively). No qualitative differences between the two infected groups were observed on HE stains in terms of granuloma formation or other inflammatory tissue changes. Infection dose 120 cercariae per mouse.

### Response to alteration of gut microbiota reflected in intestinal barrier and liver function parameters

To investigate whether the observed differences in liver pathology and lack thereof in ileum tissues were reflected in functional parameters we measured intestinal permeability and four classical clinical liver status parameters; namely, albumin (Alb), alkaline phosphatase (ALP), alanine aminotransferase (ALT) and aspartate aminotransferase (AST) (Figure 5A-E). Intestinal barrier function was assessed by FITC-dextran as a permeability measure (Figure 5F) and supported by determining expression levels of genes encoding mucins, tight junction proteins, and the c-type lectins RegIIIγ and RELM-β (Suppl. Figure S6). Both infected groups showed comparable responses to eggs traversing the intestinal barrier in terms of upregulation of expression of mucins (MUC1 and MUC2) and RELM-β encoding genes. Antibiotics administration lowered RELM-β and RegIIIγ expression. However, permeability was similar if not slightly decreased for AbxSm animals compared to infected only (p=0.078, Figure 3F). Interestingly, this corresponds to a significantly lower level of *Schistosoma* DNA per gram faeces (Suppl. Figure S5G) pointing to fewer eggs successfully leaving the hosts with altered gut microbiota. However, observations of in general larger quantities of faecal matter being passed by antibiotics-treated animals could also be the explanation for the lower *Schistosoma* DNA levels per gram faeces in the AbxSm animals.

**Figure 5:**
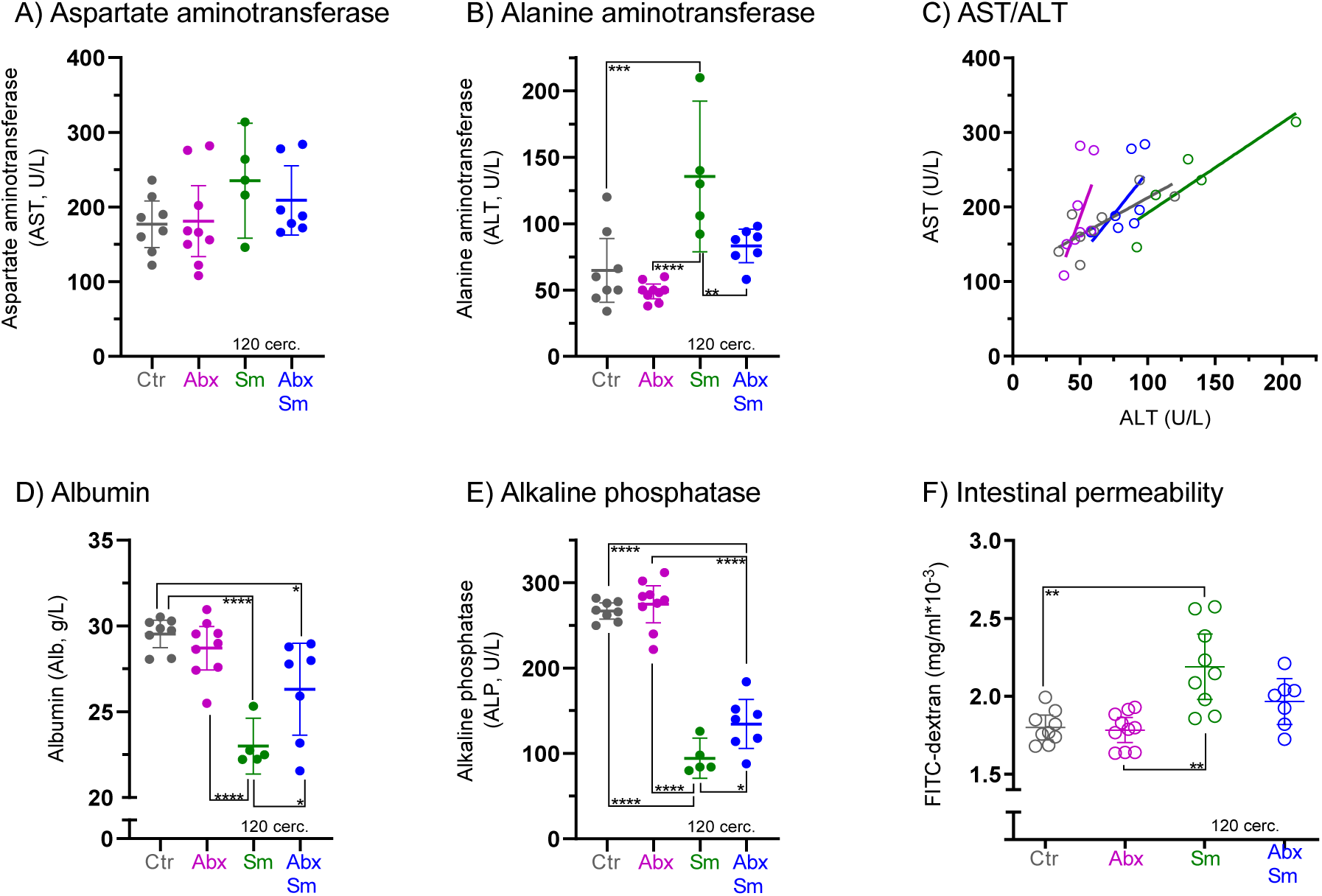
Liver function and intestinal permeability in *S. mansoni-*infected animals as affected by alteration of gut microbial composition. Serum measurements at six weeks post infection reflecting liver function and intestinal permeability (120 cercariae/infected mouse). The transaminase levels of A) aspartate aminotransferase (AST) and B) alanine aminotransferase (ALT: Ctr vs Sm p=0.0002; Abx vs Sm p<0.0001; Abx vs AbxSm p=0.054; Sm vs AbxSm groups p=0.074) point to differential liver responses between infected groups as indicated by their AST/ALT ratio with linear regression fitted lines in C), where only slopes for Ctr and Sm are significantly different to zero (Ctr p=0.027, Sm p=0.039).The AST/ALT ratios were significantly different for Ctr vs Sm (p=0.043); Abx vs Sm (p=0.0008) and Abx vs AbxSm (p=0.030). D) albumin (Alb: Ctr vs Sm p<0.0001; Ctr vs AbxSm p=0.013: Abx vs Sm p<0.0001; Abx vs AbxSm p=0.073; Sm vs AbxSm p=0.029) and E) alkaline phosphatase (ALP: Ctr vs Sm p<0.0001; Ctr vs AbxSm p<0.0001; Abx vs Sm p<0.0001; Abx vs AbxSm p<0.0001; Sm vs AbxSm p=0.040) serum levels for control (Ctr n=8), antibiotics-treated (Abx n=9), *S. mansoni*-infected (Sm n=5) and antibiotics-treated and *S. mansoni*-infected (AbxSm n=7). F) shows the relative intestinal permeability measured by FITC-dextran in serum (Ctr n=9, Abx n=10, Sm n=9, AbxSm n=7). On all plots each dot represents an individual mouse; sera were prioritised for permeability assay and then liver panel measurement, which account for the differences in n. All comparisons with adjusted p<0.1 are stated (ANOVA, Tukey’s multiple comparison test). Bars indicate means with 95% CI.

In general, we observed that liver function was clearly affected by the infection (Figure 5A-E). Elevated levels of alanine aminotransferase and albumin in the *S. mansoni-*infected groups compared to non-infected groups were observed reflecting increased liver stress. In contrast, the infected mice presented with lower levels of alkaline phosphatase compared to non-infected groups, which is consistent with the parasites consuming micronutrients. Interestingly, the AbxSm mice with altered gut microbiota show a lower level of alanine aminotransferase but a higher level of alkaline phosphatase and albumin compared to the infected-only pointing to differential response to eggs in liver associated with microbiota-mediated immune homeostasis (Figure 5B, D, E). These observations correspond to our finding of a reduced degree of inflammation in the liver in the AbxSm mice compared to the solely infected ones.

### Altered microbiota affects the host immune response induced by *S. mansoni* infection

Typical immune responses to helminth infection, such as high granulocyte numbers, more activated CD4+ cells and upregulation of MHC-II in spleens as well as cytokine production (IL-10 and IFN-y) were observed at the cellular level by flow cytometry of cells from spleens (SPL) and mesenteric lymph nodes (MLN) (Suppl. Figure S7). The AbxSm group did not present with any apparent impairment in immune response compared to the Sm group. However, a tendency towards reduced capacity of CD4+ lymphocytes to produce IFN-y in AbxSm spleens compared to Sm spleens and fewer CD103+ as well as CD69+ of FOXP3+CD4+ cells in MLNs at six weeks post infection were observed (Figure 6 AB, E). This tendency of less CD103+FOXP3+CD4+ was seen in AbxSm compared to Sm in the MLN but not the spleen, which could indicate different mechanisms behind the induction of intraepithelial Treg populations (αβ, γδ). Six weeks post infection is too early to probe whether a delay or lack of Th1 to Th2 balance shift as well as regulatory response initiation could be linked to the observed immunopathological differences between Sm and AbxSm. Therefore, another C57BL/6-NTAC model with sampling eight weeks post infection was established to profile the consequences of altered gut microbial composition on host immune homeostasis during *S. mansoni* infection on immune markers at the transcriptional level in ileum and liver and expressed level in plasma. The relative gene expression levels of a panel of immune related factors were determined in the ileum and liver tissues (Figure 6F-H, panel targets listed in legend). Principal component analysis showed a clear effect of both antibiotics treatment and *S. mansoni* infection on ileum tissue expression levels of immune system related genes (Figure 6FG). These differences were driven by multiple factors (univariate graphs shown in Suppl. Figure S8). Firstly, significantly elevated expression of transcription factors GATA-3 and FOXP3 were observed in Sm compared to all other groups. Furthermore, higher cytokine mRNA levels from IL-10, IL-6, IL-33 and IL-1β were also associated with the Sm group, which was also the case for the IL-33 receptor ST2 and the T-cell co-inhibitory CTLA-4 (Suppl. Figure S8). No differences between Sm and AbxSm mice were seen for IL-4, IL-5, IL-12 or IL-13. The antibiotics treatment combined with infection also affected expression levels of CD8, GZMB, Prf1, CXCR6 and CXCL-16 whereas TNF-α expression was affected by the antibiotics treatment only (Suppl. Figure S8). Macrophage M1/M2 phenotype markers Arg1 and NOS2 also showed differential expression patterns across the groups. Of interest, Arg1 trended towards lower expression in AbxSm compared to Sm, but both the infected groups had higher levels than controls. NOS2 was significantly lower expressed in antibiotics-treated mice compared to both Ctr and Sm, even noting a synergistic effect of infection and antibiotics treatment (Suppl. Figure S8). In contrast to at least six measured genes being differentially expressed in ileum tissue, only NOS2, IL-33 and IFN-y were significantly different in Sm compared to AbxSm mouse livers (Figure 6IJ and Suppl. Figure S9). To investigate whether these profound differences in immune response at mRNA level particularly in the ileum could be detected in circulation via a panel of immune parameters measured in plasma by a magnetic bead-based Luminex assay. Schistosomiasis (Sm/AbxSm) was clearly associated with increased levels of cytokines IL-5, IL-13, IL-17, IL-23 and GM-CSF as well as the VEGF growth factor and tissue inhibitor of MMP (TIMP)-1 compared to uninfected animals (Ctr/Abx) (Figure 6K, Suppl. Figure S10). However, no significantly different immune response between the Sm and AbxSm groups was found measuring a panel of 13 different cytokines, GM-CSF and the matrix metalloproteinases (MMP)-8, TIMP-1, and VEGF (Figure 6L). Only IL-4 levels were significantly different between the two infected groups (p=0.02).

**Figure 6:**
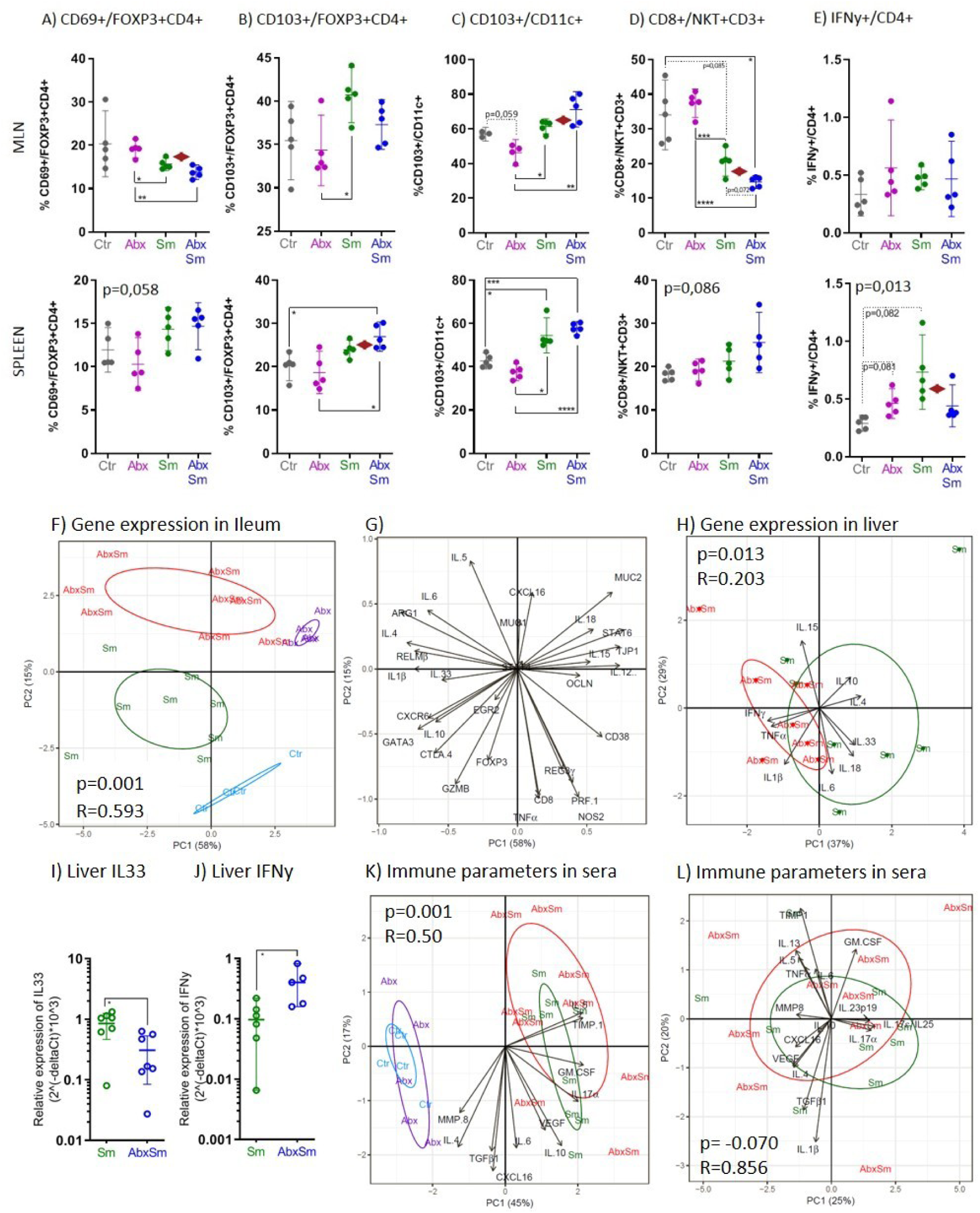
Alteration of gut microbial composition changes host immune response to *S. mansoni* infection in several tissues at six and eight weeks post infection. A-E) Immune cell phenotype proportions in mesenteric lymph nodes (MLN, top row) and spleens (second row) by flow cytometry in control (Ctr, n=5), antibiotics-treated (Abx, n=5), *S. mansoni*-infected (Sm, n=4) and antibiotics-treated *S. mansoni*-infected (AbxSm, n=5) mice six weeks post infection (120 cercariae/mouse). A) CD69+/FOXP3+CD4+, B) CD103+/FOXP3+CD4+, C) CD103+/CD11c+, D) CD8+/NKT+CD3+, E) IFNγ+/CD4+ (also see Suppl. Figure 7). Bars represent means with 95% CI. Stars indicate statistically significant differences between groups (Welch’s ANOVA with Dunnet’s correction for multiple testing, p<0,05 cut-off). A p value in the upper left corner means a significant difference detected by Welch’s ANOVA, but not by underlying pairwise testing. Red rhombes indicate p values <0.07 between Sm and AbxSm groups (t-test, Welch’s correction) for A) MLN p=0.051, B) spleen p=0.059, C) MLN p=0.055, D) MLN p=0.014 and E) spleen p=0.069. To confirm that the host immune response in Sm and AbxSm animals was truly different, gene expression profiles of ileum and liver tissues and in sera at eight weeks post infection for Ctr (n=4), Abx (n=5), Sm (n=8) and AbxSm (n=8) was obtained by Fluigdigm RT-PCR using Taqman primer-probe sets and Luminex (50 cercariae/mouse). PCA plots of the relative gene expression in ileum tissue shown in F) scores and G) loadings. Multivariate ANOSIM analyses p and R values are given on the scores plot. Sm is statistically dissimilar to all other groups F) Sm vs Ctr (p=0.009; R= 0.522), Sm vs Abx (p=0.002; R= 0,833), Sm vs AbxSm (p=0.001; R= 0.378).The two control groups are also statistically different, Ctr vs Abx (p=0.005; R=0.875). Panel targets: MUC1, MUC2, RELMβ, TJP1, OCLN, REG3γ, FOXP3, STAT4, STAT6, GATA3, EGR2, ARG1, CD38, NOS2, TNFα, IL-1β, IL-4, IL-5, IL-6, IL-10, IL-12, IL-15, IL-18, IL-33, CXCL16, CXCR6, PRF-1, GZMB, CD8, CTLA-4 (see also Suppl Figure S8). PCA biplot of relative gene expression in liver tissue from the Sm and AbxSm mice is shown in H). The liver panel consisted of cytokines (IFN-γ, TNF-α, IL-1β, IL-4, IL-6, IL-10, IL-15, IL-18, IL-33). ANOSIM (p=0.013, R=0.203). Only IFN-γ and IL-33 are significantly different between the two infected groups on a single parameter level as shown in I) and J) (see Suppl. Figure S9). K) PCA bi-plot showing scores and loadings of a panel of serum parameters with immune response relevance measured by Luminex (IL-4, IL-5, IL-6, IL-10, IL-17α, TGFβ-1, CXCL-16, GM-CSF, MMP-8, TIMP-1 and VEGF). ANOSIM p and R values are given on the biplot. The two infected groups Sm/AbxSm are statistically dissimilar to both control groups (Ctr/Abx). Ctr vs Sm (p=0.004; R=0.97), Abx vs Sm (p=0.001; R=0.95). L) The effect of antibiotics treatment between infected groups (Sm vs AbxSm) cannot be observed in circulation based on the panel except for in IL4 levels (p=0.020) as can be found in Suppl. Figure 10.

### Short-chain fatty acid (SCFA) level changes through antibiotics treatment and infection

Changes in gut microbiome composition will influence microbial metabolism and the metabolites secreted which in turn might confer a differential immune metabolic potential. Therefore, we measured SCFA levels in plasma and caecum content. SCFA levels in caecal content were as expected generally lowered in antibiotics-treated mice compared to non-treated (Figure 7A-H). However, infection (AbxSm) in addition to antibiotics treatment resulted in significantly higher propionate, butyrate, isobutyrate, 2-methylbutyrate and isovalerate levels compared to antibiotics treatment alone (Abx). A similar association as between *S. mansoni* infection and elevated SCFA levels was also observed for the branched-chain fatty acids (BCFA) isovalerate, isobutyrate and 2-methylbutyrate in infected (Sm) compared to control (Ctr) animals. In contrast, infection resulted in lower isocaproate levels both in antibiotics treated (AbxSm) and non-treated (Sm) mice compared to the control group (Ctr).

**Figure 7:**
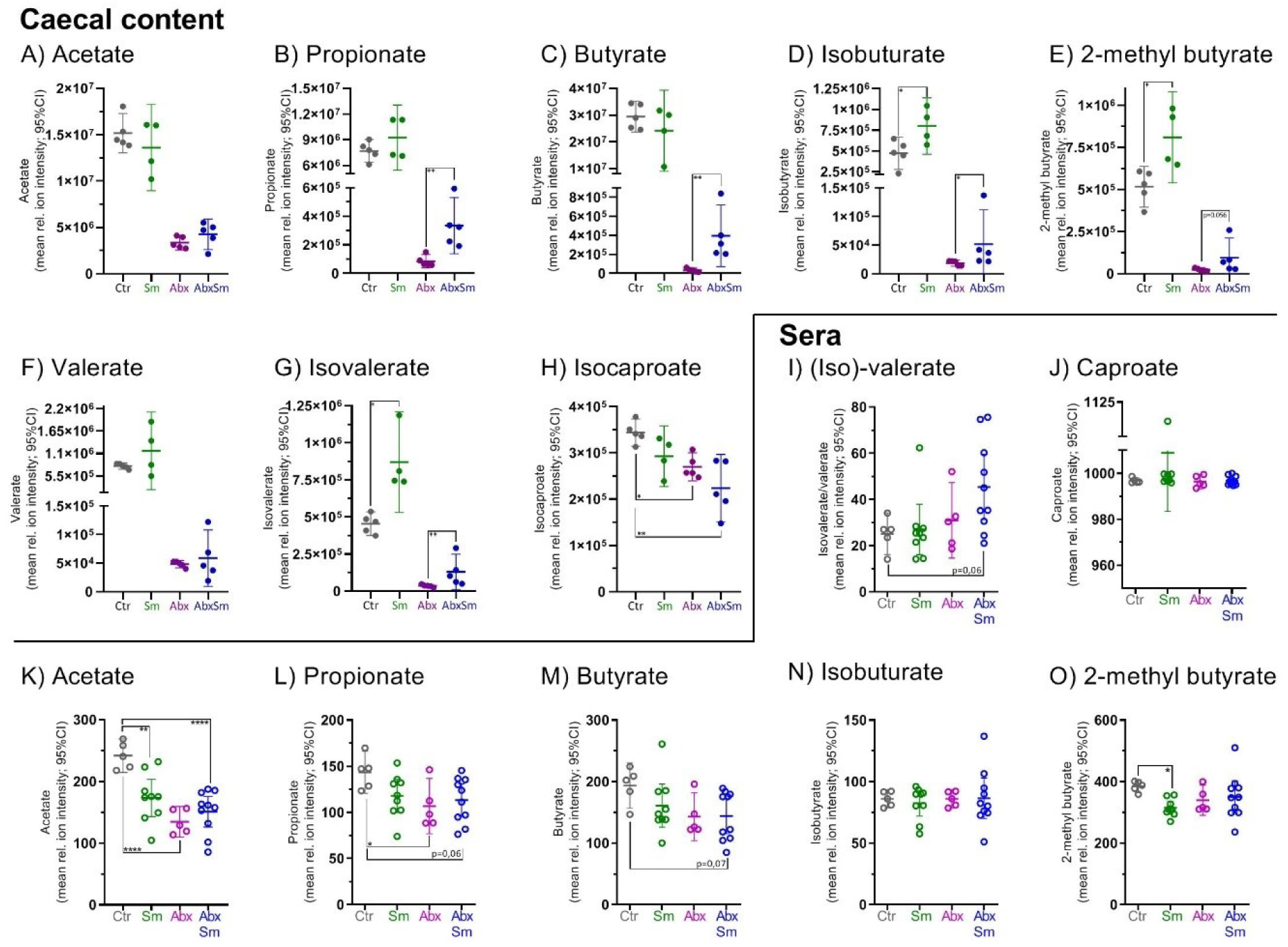
SCFA levels in sera and caecal content changes in response to *S. mansoni* infection. SCFA levels measured by LC-MS in A-H) caecal content and I-O) sera. SCFA levels in caecal content for A) acetate, B) propionate, C) butyrate, D) isobutyrate, E) 2-methyl butyrate, F) valerate, G) isovalerate and H) isocaproate from control (Ctr, n=5), antibiotics-treated (Abx, n=5), *S. mansoni-* infected (Sm, n=4) and antibiotics-treated *S. mansoni-*infected (AbxSm, n=5) mice eight weeks post infection with 50 cercariae. SCFA levels in sera is shown in I) (iso)valerate, J) caproate, K) acetate, L) propionate, M) buturate, N) isobutyrate and O) 2-methyl butyrate for Ctr (n=5), Abx (n=5), Sm (n=9) and AbxSm (n=10) mice at eight weeks post infection. Significant differences determined by ANOVA with Dunnet’s multiple comparisons test are indicated with stars. Adjusted p values under 0,1 are given on the graphs. Antibiotics treatment clearly significantly reduces SCFA levels in caecal content, which is not indicated on graphs (A-H).

While caecal content SCFA levels can impact local gut immune responses, we considered whether SCFA levels in the systemic circulation also would be altered as a result of antibiotics treatment and parasite infection. Antibiotics treatment (Abx) conferred lower SCFA levels in plasma for acetate, propionate and butyrate despite lower levels for all measured SCFAs in caecal content compared to controls (Ctr). In general, plasma levels of SCFAs were slightly lower in infected (Sm) compared to control (Ctr) animals (Figure 7I-O). The most pronounced differences were observed for acetate and 2-methylbutyrate. For AbxSm and Abx mice SCFA levels were comparable.

## Discussion

In our C57BL/6-NTac mouse model, the effects of *S. mansoni* infection on gut microbial composition are driven by worms (prepatent) and additionally by eggs (patent) even during antibiotics treatment. The findings confirm prior reports of *S. mansoni* inducing gut microbial composition changes in mouse models (25, 29, 30). However, the compositional changes differ from study to study which is likely due to differences in local housing milieu, age and mouse strain used, as well as basic experimental set-up differences. For example, Floudas *et al.* (2019) found that a higher relative abundance of *Alistipes* sp. being associated with infection (25, 29), while we observe the opposite. The presence of worms alone affecting faecal microbiome composition is likely initiated by immune processes started by cercarial penetration of skin and schistosomula migrating through lung capillaries causing activation of the gut mucosal immune compartment (31, 32). The worms themselves may also alter the gut microbial milieu e.g. via their own metabolism and/or activities of the microbes they harbour (72). Infection-driven local immune environment changes facilitate subsequent differential colonisation; for example an increased mucin production favour mucin-degrading species as seen in our model by upregulation of MUC1 and MUC2 which corresponds to a higher relative abundance of *Bacteroides* spp. in infected compared to uninfected animals (33). Infection, and in particular, patent infection was clearly linked to the alteration of faecal microbiome composition. However, the alterations of community composition at the individual level are complex and liable to differential colonisation rather than a consistent infection-driven signature composition. These observations are consistent with the limited and heterogenous faecal microbiome differences associated with both *S. mansoni* and *S. haematobium* infection in children and adolescents (34–37). Complex environmental as well as host factors influence infection-associated gut microbial community changes warranting caution when extrapolating animal model microbiome composition to human infection context (38, 39). As *S. mansoni* infection influenced gut microbial composition, the question was whether an altered gut microbial community structure prior to infection would in turn impact immunopathology. A previous study by Holzscheiter *et al.* (2014) investigated immunopathological consequences of short-term microbiota depletion using a cocktail of antibacterial, –fungal, and –protozoal drugs in mice with already established *S. mansoni* infection, wherein no effect on liver granulomas but reduced intestinal inflammation and granuloma formation were observed (18). In contrast, our model is based on host immune responses to primary *S. mansoni* infection in mice with prior and upkept ampicillin and vancomycin mediated alteration rather than depletion of the gut microbial community composition. Using stereology, we demonstrate immunopathological consequences in the liver but not the ileum of *S. mansoni*-infected mice with altered gut microbial composition (AbxSm) compared to mice with unmanipulated gut microbiota (Sm). The *S. mansoni*-infected groups showed a similar degree of hepatomegaly, but the relative degree of inflammation in liver tissue was decreased in the AbxSm animals. This difference could not be explained by infection burden alone neither in terms of eggs, worm numbers, or sex ratio (40), but is consistent with the observed higher liver IFN-ɣ expression level in the AbxSm group at eight weeks post infection (41). IFN-ɣ is typically associated with the prepatent and early patent phase of *S. mansoni* infection where a strong Th2 response still has not been initiated (42). In the Sm group significantly higher expression levels of IL-33 and IL-4 in blood were found, which is indicative of a Th2 polarisation and granuloma formation (43–45). Notably, comparable levels of the major egg-secreted antigens IPSE/alpha-1 and omega-1 were seen in both groups (10, 46). The comparable infection intensities and excretory/secretory capacity of the eggs point to the antibiotics administration having no direct effects on the parasites. Albumin, alanine aminotransferase (ALT), and alkaline phosphatase (ALP) levels further support that the inflammatory liver environments in Sm and AbxSm animals were different six weeks post infection. Elevated levels of ALP are frequently seen in response to infectious conditions and liver disease such as hepatitis or cirrhosis, including *S. mansoni* infections. This is in contrast to the current study, where lower levels were observed (47–49). However, low values have been associated with protein malnutrition, and vitamin and mineral deficiencies (50, 51). This is in line with effects conferred by the relatively high burden of adult worms’ blood feeding in relatively small host animals (Suppl. Figure S5A), despite the *ad libitum* access to food, and potentially combined with altered absorption capacity due to inflammatory responses initiated by eggs traversing the intestinal wall. Intestinal barrier function affects the number of bacteria and bacterial epitopes translocating to circulation and hence to the liver, where they can enhance inflammation (52, 53). As we see a trend towards lower permeability in the AbxSm animal compared to Sm only, it is possible that bacterial translocation is changed in numbers and nature due to a restricted number of species surviving the antibiotics. A previous study implicates a role for certain organisms in the gut commensal milieu in *S. mansoni*-induced pathology by showing less DNA damage and hence pathology in association with administration of the probiotic *Weizmania coagulans* (54, 55). We observed no difference in the degree of inflammation in the ileum, which was somewhat surprising. It is possible that the *S. mansoni* egg stimuli during the up to two weeks of patent infection before termination as well as the remaining gut microbiota provide sufficient stimuli for “normal” egg passage. Consistent with Bloch (1980), we saw both passage of single/few eggs and clustered rows migrating across the intestinal barrier always moving parallel to the lumen of the intestine before travelling towards the lumen (7). This is in line with previous studies demonstrating that eggs exploit and induce vasculature in association with a predilection for Peyer’s patches (56). We observed no differences related to the antibiotics treatment with respect to egg migration, which varied from none to major inflammation and granuloma formation. Despite no apparent effect on infection burden, the fecundity of worms or egg viability, we observed consistently fewer eggs and/or DNA in faeces for mice receiving antibiotics, which was also reported by Holzscheiter *et al.* (18). Taken together our observations regarding the liver showed less granulomatous tissue, fewer granulomas with minor collagen depositions, altered liver enzyme levels, and local Th1/Th2 milieu differences indicated by expression of more IFN-ɣ and less IL-33 in the AbxSm mice compared to the Sm group. This suggests, that the immune responses induced by *S. mansoni* eggs are delayed or attenuated in the livers of mice with altered gut microbiota (AbxSm). Furthermore, patent *S. mansoni* infection, whether under antibiotic pressure or not, drives changes in SCFA levels, gut microbiome composition as well as distinct immune milieu changes in local and systemic contexts.

This study is the first experimental schistosomiasis model addressing both how the infection can drive a change in gut microbial composition combined with how an altered gut microbiome in turn can affect host immune responses and immunopathology related to infection with the parasite. In general, our results support a pivotal role of intestinal immune homeostasis in the regulation of systemic pathology in *S. mansoni* infection mediated by intertwined host-parasite-commensal gut microbiota interactions. It is tempting to speculate that more subtle alterations of gut microbial compositions mediated by factors such as diet, pre– and probiotics, other infections, transient antibiotic intake, or other drugs also can facilitate changes in the immune milieu that affects systemic pathology. In a translational context, future human studies could benefit from longitudinal cohort designs when attempting to unravel to what extend and under which conditions certain gut microbial communities and metabolic signatures may impact immunopathology dynamics and morbidity; and vice versa in terms of understanding which infection pressures and durations of *S. mansoni* infections drive persistent changes intestinal immune milieus and gut microbiota composition.

## Materials and methods

### Ethics

Mouse experiments were performed according to the Danish Act on Animal Experimentation (LBK no. 474 of 15/05/2014). The study was approved by the Animal Experimentation Inspectorate, Ministry of Environment and Food, Denmark (license no. 2015-15-0201-00694). Health monitoring was done according to FELASA guidelines (57).

### Animal model

Female C57BL/6-NTac mice (Taconic, Denmark) were housed in open-lid cages with enrichment in our AAALAC accredited barrier-protected facility on 12:12 hour light:dark cycle from six weeks of age. The mice were fed Altromin 1324 *ad libitum* (Brogaarden, Denmark) and tap water. All experiments were based on a study design with four groups: control (Ctr), antibiotics control (Abx), *S. mansoni*-infected (Sm), and antibiotics-treated + *S. mansoni*-infected (AbxSm). Cage and group allocation was done by a person blinded to group category. A cocktail of 1g/L ampicillin (Boehringer-Ingelheim) and 0.5g/L vancomycin (Hospira) was administered via drinking water to group Abx and AbxSm animals and renewed twice a week (waste collected and disposed of according to local authority guidelines) from day 0 at 7 weeks of age and to termination. Group Sm and AbxSm mice were percutaneously exposed to *S. mansoni* cercariae a minimum of one week after the start of the administration of antibiotics. Data from three independent experiments are used in this paper. As we observed higher than expected pathogenicity for the Puerto Rican (PR) strain in C57BL/6-NTac mice at six weeks post infection (dose 120 cercariae/mouse), the dose was reduced in the experiment which was terminated at eight weeks post infection (50 cercariae/mouse). Sample sizes vary per assay due to the death of some mice, differences in available sample volume or technical reasons; n is given per group in each figure legend. Cercariae came from the *S. mansoni* life cycle maintained locally at the Section for Parasitology and Aquatic Pathobiology, University of Copenhagen based on the *S. mansoni* PR strain provided by Mike Doenhoff in 2015 and propagated in female NMRI:BomTac mice (Taconic, Denmark) and *Biomphalaria glabrata* snails originally obtained in 2002 from the Theodor Bilharz Institute in Egypt. Infection was confirmed either by presence of circulating cathodic antigen in 20 µl mouse urine detected with the POC-CCA test (Rapid Medical Diagnostics), observations of hepatosplenomegaly and/or *Schistosoma* spp. DNA in faeces (described below).

### 16S rRNA gene amplicon based faecal and caecal microbiome characterisation

Faecal pellets were collected from individual mice placed briefly in individual clean cages. Each group consisted of animals housed in two separate cages. After collection faecal samples were stored at –80°C until further analysis. DNA was extracted using the PowerSoil DNA isolation kit (MoBio Laboratories Inc) according to the manufacturer’s protocol with the following adjustments; max 150 mg faecal pellets put directly into bead-tubes and vortexed briefly were heat-treated for 10 min at 65°C followed by 10min at 95°C, whereafter samples were cooled on ice before being subjected to 3×15 seconds 6,5M/s bead beating on a Fastprep (MP Biomedicals). A volume of 2 µl extracted DNA was measured on NanoDrop 2000 and samples were stored at –60°C. After DNA extraction all samples were given a computer-generated random ID number and analysed in that order to minimise biases introduced in the subsequent library building process. A 16S rRNA gene amplicon libraries targeting the V3 region were made as described in Krych *et al.* (2018) (58) and sequenced on a NextSeq (Illumina, USA) platform using MID 2×150 chemistry. Sequences were demultiplexed by Perl script, (59) and run through the IMNGS platform (60) based on the UPARSE algorithm (61) with an abundance cut-off of 0.001. The obtained sequences were run through the SILVA aligner (62, 63) for improved taxonomy annotation. If taxonomy was not obtained through SILVA, but a classification was assigned by IMNGS, the IMNGS annotation was kept. OTUs where the total number of reads was ≤100 across all samples were excluded. Negative controls were included in all steps from DNA extraction and onwards and sequenced and used to identify potential contaminants. Sequences were re-aligned and a neighbor joining tree (bootstrap, maximum composite likelihood, pairwise deletion) was generated using MUSCLE alignment in MEGA7 (64). Data was then processed using the RHEA package pipeline (65) using R with cumulative sum scaling as normalisation (66, 67). The Shannon effective index was used to describe α-diversity and a generalised UniFrac model for β-diversity with non-metric multidimensional scaling plots and PERMANOVA analysis (68).

### *S. mansoni* DNA in faeces

DNA was extracted as described above for 16S rRNA gene amplicon sequencing. *Schistosoma* spp. internal transcribed spacer 2 (ITS2) specific primers as published by Obeng *et al.* (69) were theoretically optimised based on the *S. mansoni* ITS2 sequence (GenBank sequence AF503487.1) to the following Sm48F 5’-GGTCTAGATGACTTGAT<u>T</u>GAGATGCT-3’ and Sm124R 5’-TCCCGAGCG<u>T</u>GTATAATGTCATTA-3’. No sequence change was done for the Sm78T probe (5’-FAM-TGGGTTGTGCTCGAGTCGTGGC-BHQ1-3’). Semi-quantitative qPCR was run on the Bio-Rad CFX Maestro 1.0 platform (3 min hot-start (95⁰C); 45 cycles; 15 sec 95⁰C, 30 sec 60⁰C) with DreamTaq Hot Start PCR Master Mix (2x) (ThermoFischer), primers (4 µM)/probe (2 µM) and 10 ng extracted faecal DNA/20 µl reaction in duplicates with a known positive control of *S. mansoni* DNA extracted from worms (Qiagen DNeasy Blood and Tissue Kit) and negative control faecal DNA from uninfected mice.

### Liver digests

Partial livers from Sm and SmAbx mice (eight weeks post infection) stored at –20⁰C were thawed, weighed and digested at room temperature in 40 ml 4% KOH on magnetic stirrers. Digestion was slowed by adding 40 ml H_2_O. After mixing thoroughly six one ml subsamples were counted in Sedgewick-Rafter chambers with glass coverslips at 10x magnification by two blinded investigators (three samples each from each digest per investigator). The number of eggs was calculated to eggs/gram liver tissue.

### Volume and degree of inflammation quantified by stereology

Entire livers from each of the four experimental groups were fixed in 10% neutral buffered formalin (NBF, Sigma), dehydrated in increasing alcohol concentrations, and embedded in paraffin (LeicaASP300 tissue processor) before being sliced in parallel 8 µm thick sections on a Jung sledge microtome. Every 40^th^ section was mounted on Superfrost Plus microscope slides (Fisherbrand). The first section to be sampled was determined from a random number table for each liver, choosing a random number within the sampling frequency (in the liver; k=40) and continuing systematically in a predetermined manner, for example section 13-53-93 etc. Sections were dried and then heated to 60°C for 40 minutes, dewaxed in xylene and a series of dilutions of ethanol (99, 96, 70%) and subsequently stained with standard HE stained (Mayer’s haematoxylin 6 min, 1% eosin Y (Sigma) 4 min). The liver volume was estimated using the Visiopharm newCast system (6.5.0.2303) where software generated systematic uniform random sampling (SURS) at 4x magnification. A point count was done using Cavalierís method for volume estimation as described for *S. mansoni* infected livers in Friis *et al.* (1998) (27, 28). Briefly, for each point on a point-counting grid, there is a corresponding area, a(p), which is a constant. Counting the total number of points hitting the object of interest, ∑P, and knowing the area associated with each point the area of the region of interest (ROI) can be obtained per sampled section. The volume of the ROI is equal to V(object) = t x k x a(p) x ∑P (28) where (t) is the section thickness = 8 µm, and the sum of P is the total number of points counted on all sampled sections. For livers the area per point were 9mm^2^ (group Ctr and Abx) and 20.25mm^2^ (group Sm and AbxSm), respectively. For Ctr/Abx livers 141-206 points were counted (13-15 sections/liver) and the range for Sm/AbxSm livers was 130-187 points (13-18 sections/liver). A relative measure for the proportion of inflamed tissue in the livers was obtained by also counting every time point(s) with an a(p)=0.07mm^2^ in a 3×3 grid (0.65 mm^2^) inside a defined counting frame including the central point used for liver volume estimation would hit on granulomatous tissue (suppl. Figure S3). Furthermore, the number of eggs observed in each counting frame was recorded and a count of the number of worm cross-sections on all liver was sections performed. Tissue processing, staining and systematic random sampling for volume and relative degree of inflammation were also done on ileum sections as described for livers with the following adjustments. Three consecutive one-centimeter-long pieces from the caecum and in the proximal direction (ileum, ileum, ileum/jejunum) were embedded in paraffin within a vertical axis. This resulted in three cross-sections of ileum tissue on each slide. Every 60^th^ 8 µm thick parallel section cut on a Leica RM2125 RTS microtome was sampled until a total of twelve sections was obtained from each animal (36 cross sections/animal). For ilea the area per point counted was 0.09 mm^2^ and 0.04 mm^2^ in a 3×3 grid for the proportion of inflammation for all groups and the count was done at 10x magnification. For ileum cross sections a minimum to a maximum of 352-438 for Ctr, Abx and of 537-1431 for Sm, AbxSm sampling points were counted due to large biological variation. “Inflammation” in ileum sections was defined as a point hitting either granuloma, lymphocyte-rich cell infiltrates, clear remodelling of and/or cell infiltration in tissue (e.g. markedly changed submucosa after egg passage). Hence, the defined term of “inflammation” in the ilea consists of a range of deviations from normal tissue. The number of eggs visible at 10x magnification in each entire ileum cross-section was recorded. For both the liver and the ileum points hitting eggs were counted as “inflammation”. All counts on both liver and ileum tissue were done fully blinded by the same investigator.

### Histology

Immunohistological staining of the secreted egg antigens IPSE/alpha-1 and omega-1 on liver and ileum sections was performed with the mouse monoclonal antibodies anti-IPSE/alpha-1 (clone 74 1G2) (70) and anti-omega-1 (140 3E11) (10), respectively. Both antibodies were diluted 1:100. Antibody binding was visualized using the M.O.M. ImmPRESS HRP Polymer Kit and the substrate Vector NovaRed (Vector Laboratories) according to the manufacturer’s instructions. Pictures were taken with the Olympus BX51 microscope. Mayer’s haematoxylin (Merck) was used for counter staining.

Standard Masson’s trichrome stain was done on the liver section from *S. mansoni*-infected mice. Qualitative observations were made in a blinded fashion for collagen deposition which were categorised as “minor” or “major” and counted separately in association with granulomas and small vessels for one large liver section/animal by light microscopy at 10x magnification using DinoCapture 2.0 software and Dino-Eye microscope eye-piece camera (AM423x). The area of the liver section was estimated to be an average of two adjacent sections from the stereology dataset.

### Intestinal permeability and liver enzymes

Mice were fasted for four hours prior to oral gavage of 0.6 mg/g body weight FITC-dextran (4 kDa, FD-4 Sigma-Aldrich) dissolved in PBS. Dosage was done at two-minute intervals between the mice and after two hours the mice were bled via the retro-orbital vein using sodium-heparin-coated micro-haematocrit capillary tubes (Brand) under full anaesthesia (Hypnorm (Vetapharm, UK) / Midazolam (B. Braun Medical, Denmark)) and subsequently euthanized by cervical dislocation in the same order as gavaging took place. Blood was spun at 4°C for 7 min (8000xg), and plasma was kept dark and cold until analysis. Care was taken to avoid haemolysis and a subsample was aliquoted for liver enzymes measurements (stored at –20⁰C until use). Duplicate 1:1 dilutions of plasma to PBS were measured against a two-fold dilution array standard curve of (8-0.125 µg/ml) FITC-dextran in fresh plasma (1:1 in PBS) by spectrophoto-fluorometry (SpectraMAX GeminiXS) with an excitation of 485 nm and an emission wavelength of 535 nm in a 96-well NUNC flat bottom plate. To determine liver enzyme and albumin levels in plasma samples, they were thawed and diluted 1:1 in sterile H_2_O and the contents of albumin (Alb), alanine aminotransferase (ALT), aspartate aminotransferase (AST) and alkaline phosphatase (ALP) were measured on the Advia 1800 Clinical Chemistry System (Siemens).

### Flow cytometry

Cells were isolated from mesenteric lymph nodes (MLN) and spleens by mechanical disruption of the organs, filtration to remove large tissue residues (BD filter), ACK-buffer red blood cells lysis for spleens, centrifugation, and resuspension in cold PBS (2% FCS). For staining the following goat anti-mouse antibody combinations for MLN and spleen cells were used 1) surface αCD103-FITc,αCD4-PerCPcy5.5, αCD69-APC intracellular αFOXP3-PE, 2) surface αCD3-FITc, αNK/NKT-PE (eBiosciences), αCD8a-PerCPcy5.5, 3) surface αCD11c-FITc, αCD103-PE, αF4/80-PerCP_cy5.5, αMHCII-APC, 4) surface αCD8-PerCPcy5.5, αCD4-APC intracellular αIL10-FITc, αIFN-y-PE. Cells for 4) were stimulated at 37°C for 4-5 hrs with PMA (50ng/ml) and ionomycin (750ng/ml) in the presence of Golgi-stop (0,67 µl/ml) in RPMI prior to staining. After surface staining, cells for 1) and 4) were fixed and permeabilised for intracellular staining. A BD Accuri C6 Cytometer was used to acquire cells and FlowJo used to process data.

### Gene expression

RNA was extracted from an ileum and a liver piece and stored in RNAlater (Sigma-Aldrich) by homogenising the samples in 400 µl lysis buffer with 2.8 µl β-mercaptoethanol and acid-washed glass beads (Sigma-Adrich) on a FastPrep-24 (6.5m/s, 45 seconds x2) (MP Biomedicals) and subsequent processing with MagMAX™-96 Total RNA Isolation Kit (AM1830, Life Technologies) according to manufacturer’s protocol. The isolated RNA was then DNAase treated with TURBO DNA-free Kit ™ (Life Technologies) before cDNA was synthesized (High Capacity cDNA Reverse Transcription Kit, Applied Biosystems) both according to the manufacturer’s protocol. Expression levels of a panel of immune response relevant genes was measured by real-time PCR using exon-spanning TaqMan Gene Expression Assays (Suppl. Table 1) on the Biomark HD platform as per manufacturer’s instruction (Fluigdigm). Data was obtained with BioMark™ HD Data Collection Software v.3.0.2 and analysed using Fluidigm® Real-Time PCR Analysis Software v.4.1.3. The relative expression of target cytokines was calculated by normalising to ACTB expression (2^(-ΔCT)). Samples with ACTB cycle threshold values between 5 and 21 were accepted.

Multivariate principal component analysis (PCA) was done using R (3.5.0 Joy of Playing). Sample variables were normalized by Centered Log Ratio transformation (CLR) and scaled to unit variance before PCA analysis. To test group differences ANOSIM with euclidian distances as sample similarity measure (999 permutations) was used.

### Immune parameters measured by Luminex

Mice were retro-orbitally bled under full anaesthesia (hypnorm/midazolam) using sodium heparinate-coated micro-haematocrit capillary tubes (Brand) and euthanized by cervical dislocation. Blood samples were placed on ice, spun down at 5⁰C within 20 min and the plasma was carefully transferred to two clean tubes while avoiding hemolysis (one aliquot for SCFA measurements) plasma was snap-frozen, transported on dry ice and stored at –80⁰C until use. A custom premixed mouse multi-analyte magnetic Luminex assay (R & D systems) was performed according to the manufacturer’s protocol. Samples were run in 1:4 dilutions and read on the Luminex LX200 system with XPONENT 3.1 software (R & D systems). The panel consisted of the following analytes for all groups: TNF-α, IL-1β, IL-13, IL-17ε/IL-25, IL-23p19, IL-4, IL-5, IL-6, IL-10, IL-17α, CXCL-16, GM-CSF, MMP-8, TIMP-1 and VEGF. The magnetic bead TGF-β1 base kit was also used according to the manufacturer’s instructions (R & D systems) and run separately. Multivariate principal component analysis (PCA) was done using R.

### Short-chain fatty acids (SCFA) in caecum content and plasma

Caecum content samples were weighed and mixed with 1:10 w/v 50% aqueous acetonitrile (ACN) added (1:10 w/v). The mix was then vigorously vortexed for 1 min to extract the SCFAs followed by centrifugation (4000g, 10°C, 10 min) and the supernatant was collected and stored at –80⁰C until subsequent LC-MS measurements. Samples for SCFA measurements in plasma were obtained as described above. SCFAs were measured by LC-MS using a protocol modified from Han *et al.* (2015) (71). 20 µl of sample were transferred to glass tubes, and derivatisation was initiated by addition of 20 µl 200 mM 3-nitrophenylhydrazine (3NPH)·HCl in 5:1 water:ACN, followed by 20 µl of 120 mM *N*-(3-dimethylaminopropyl)-*N*′-ethylcarbodiimide (EDC)·HCl in 6% pyridine:water. Samples were vortexed and incubated at 40°C for 30 min, before 10 µl of the derivatised sample was transferred to pre-chilled Eppendorf tubes containing 490 µl 1:1 ACN:H_2_O with internal standard spiked in. After centrifugation at 16.000xg for 5 min, 100 µl of the diluted sample was transferred to glass vials and kept at 4°C until analysis. SCFA 3-nitrophenylhydrazones were measured on a Thermo Vanquish UHPLC System coupled to a Thermo TSQ Quantiva Triple Quadrupole Mass Spectrometer. Chromatographic separation was performed on a BEH C_18_ (2.1 × 100 mm, 1.7 μm) column, with water:formic acid (100:0.01, v/v) as solvent A, and ACN:formic acid (100:0.01, v/v) as solvent B. Flow rate was kept at 0.35 ml/min with an injection volume of 3 µl, and a binary solvent elution gradient as described in Suppl table 2. MS analysis was carried out in negative ionization mode. Peak annotation and integration were carried out in Thermo Xcalibur 2.2 SP1.48. Peak areas were normalized to internal standard areas and calculated as relative ion intensities.

### Data analyses

Unless specifically described in the above methods sections, data were analysed and graphical representations were made using R software and GraphPad Prism. Depending on the data distribution, Student’s t-tests (Welch’s correction) or Mann-Whitney (2-sample) were applied for comparisons between two groups. K-sample tests were correspondingly done with ANOVA (Tukey’s multiple correction test adjusted), Welch’s ANOVA (Dunnett’s correction for multiple testing), and Kruskall-Wallis (Dunn’s correction) statistical tests. P values below 0.05 were considered significant. Linear regression analysis was used to investigate relations between the number of eggs versus the volume and proportion of inflamed tissue.

## Acknowledgements

We are grateful to Susanne Sørensen and Katrine Fabricius for expert stereology guidance, to Claus Stjernegaard/Veterinært Diagnostisk Laboratorium and Elisabeth Wairimu Petersen for technical assistance, and to Mette Nelander and Helene Farlov for animal work assistance. We also thank Rune Stensvold (SSI) for excellent advice during the project. Lastly, a special recognition to the late Prof. Bente Pakkenberg for her invaluable guidance, dedication to science, and hospitality at Bispeberg Neuro-stereology Laboratory.

This work was only possible thanks to a PhD-scholarship from the Faculty of Health and Medical Sciences, University of Copenhagen awarded to A. Kildemoes (R45-A3350) and Lundbeck Foundation running costs grant (R183-2014-3032).

## Supplementary figures Kildemoes *et al*

**Suppl. Figure S1:**
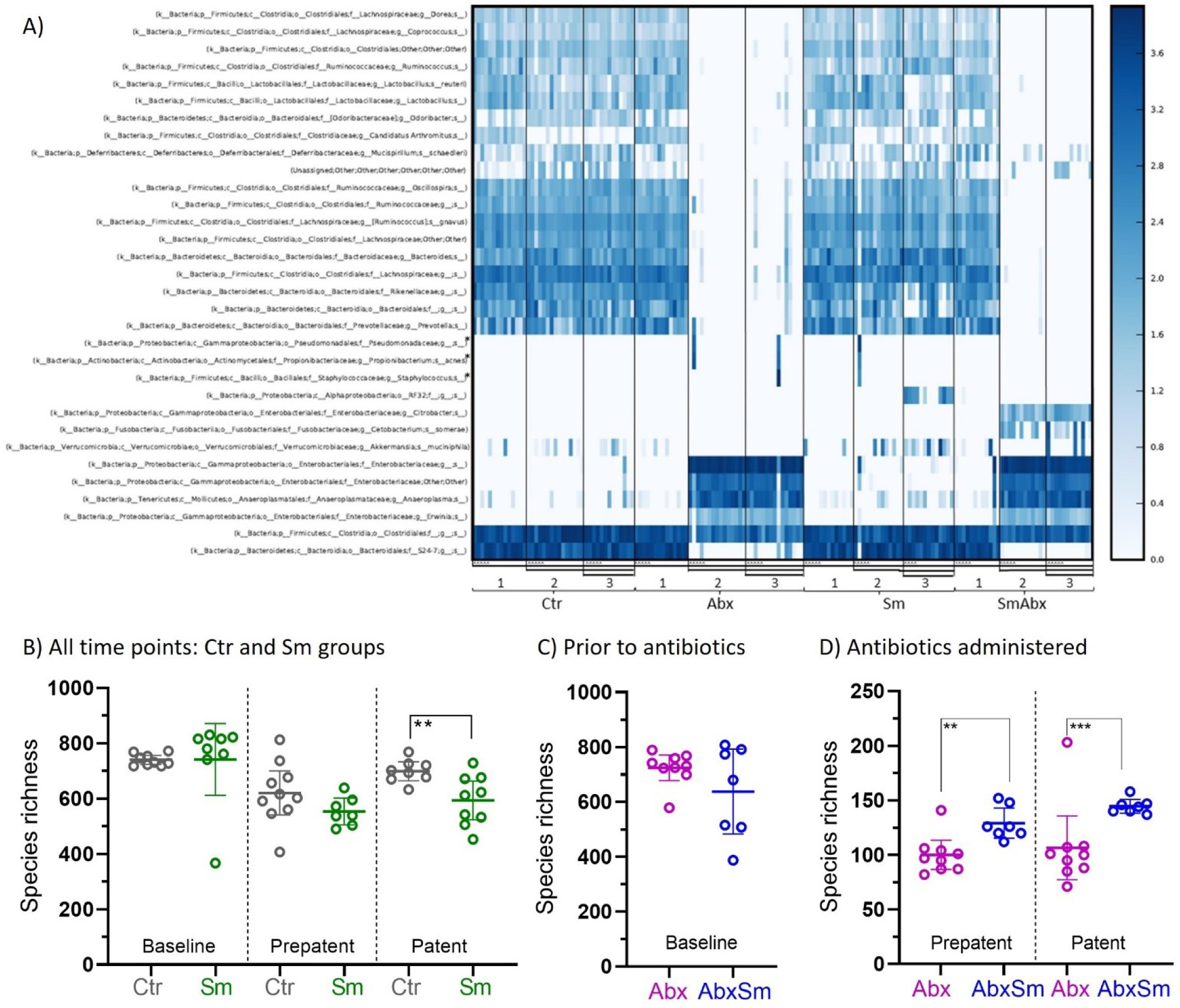
The experimental setup produces reproducible output and the antibiotics cocktail is efficacious. Heatmap of the relative frequency of OTUs (species level taxonomy) for baseline (1), prepatent (2), and patent (3) time points as indicated with horizontal lines. Here samples from an independent experiment to reproduce results presented in this paper are included (n=5 per group as indicated with ^). Total group sizes are as follows for Control (Ctr, n_total_ Time 1 = 14, Time 2 = 15, Time 3 = 13), Antibiotics treated (Abx n_total_ Time 1 = 14, Time 2 = 15, Time 3 = 15), *S. mansoni* infected (Sm n_total_ Time 1 = 13, Time 2 = 12, Time 3 = 13), *S. mansoni* infected and antibiotic treated (SmAbx n_total_ Time 1 = 12, Time 2 = 12, Time 3 = 12). *Contaminants. The heatmap illustrates the effectiveness of the antibiotics cocktail at time 2 and time 3 administered to the Abx and AbxSm groups from immediately after baseline sampling (as also seen on D). B) Species richness shown for the Ctr and Sm groups stratified by time point. C) Species richness shown for the baseline time point for the Abx and AbxSm groups, where samples were collected just prior to the administration of antibiotics. D) Species richness shown for the prepatent and patent time points for the Abx and AbxSm groups. B-D) Statistical comparison made by t-test (Welch’s correction).

**Suppl. Figure S2:**
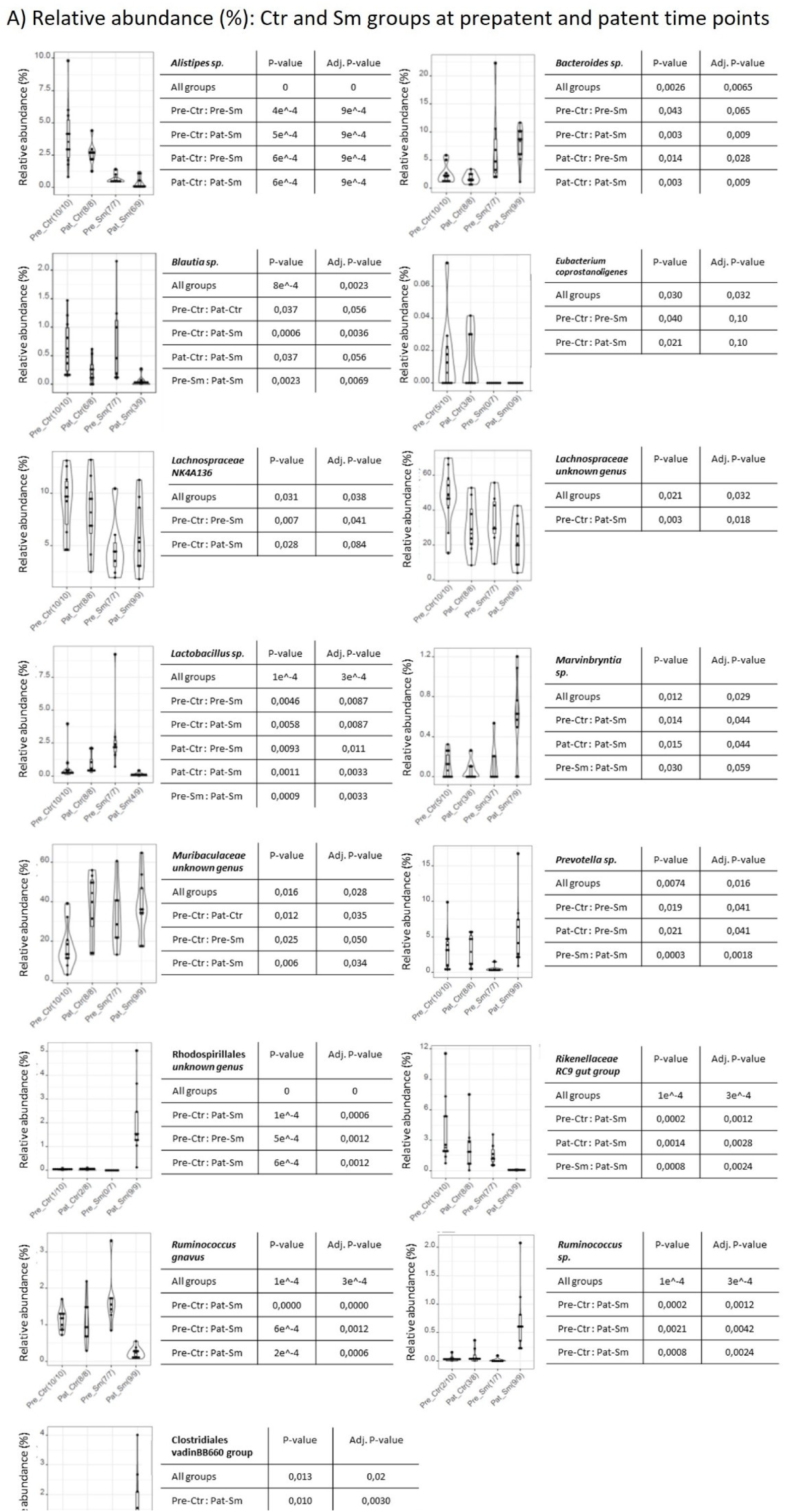
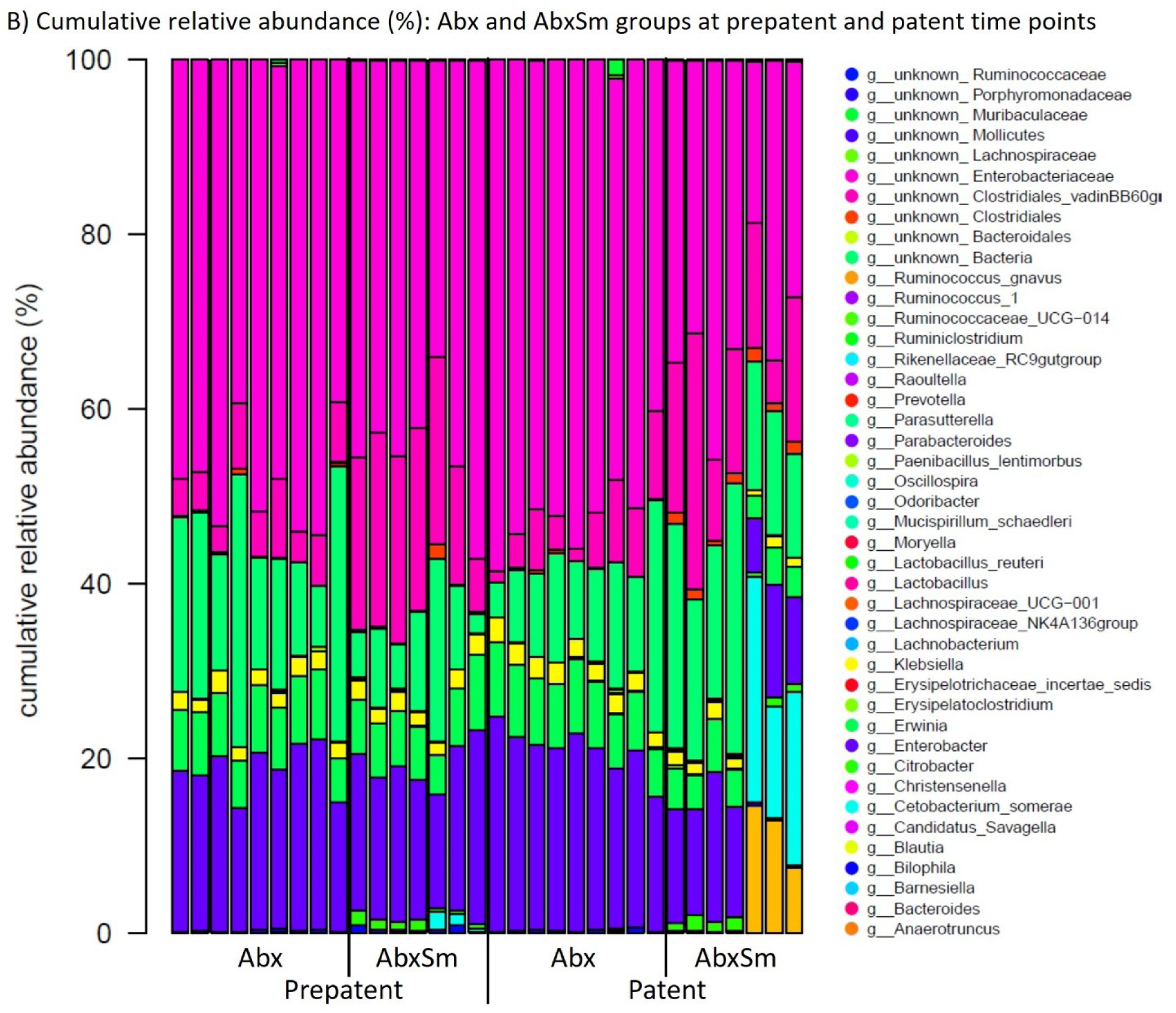
Relevant differences in 16S rRNA gene amplicon based composition at genus level at prepatent and patent time points. A) Relative abundance (%) of relevant genera for the Ctr and Sm groups at the prepatent and patent time point, where any non-parametric group comparison was significantly different at p<0,05 (analyses cut-offs: abundance 0,05 (lower not considered); prevalence 0,3 (presence of a taxonomic parameter present in at least 30% of one group); median for inclusion in testing 0,5 (very low abundant taxonomic groups not tested)). P-values unadjusted and adjusted for multiple testing (Benjamini-Hochberg method) are given for overall group differences by Kruskall-Wallis Rank Sum Test (all groups) and between groups by Wilcoxon Rank Sum Test (pairwise group comparisons; note that comparisons between groups at the same time point (pre-pre; pat-pat) or within groups over time (pre-pat for Ctr and/or Sm) are the most relevant observations). Abx and AbxSm groups were also compared, however, since the cocktail antibiotics treatment is highly efficacious, no significant differences between groups at p<0,05 were found even with the following low analyses cut-offs: abundance 0,001; prevalence 0,2; the median for inclusion in testing 0,01. B) shows a cumulative abundance plot of the prepatent and patent time points for the antibiotics treated groups. The genera/species most notably associated with infection are *Citrobacter* spp.*, Cetobacterium somerae,* and *Anaerotruncus* spp. (note the strong cage-effect at the patent AbxSm time point where the four first and three last samples came from co-housed mice). Additionally, very low abundant reads from *Alistipes* spp. *Acetatifactor* spp.*, Akkermansia municiphila, and Lachnospiraceae A2* were found and are included in the cumulative relative abundance plot, however, but are not featured on the genus list.

**Suppl. Figure S3:**
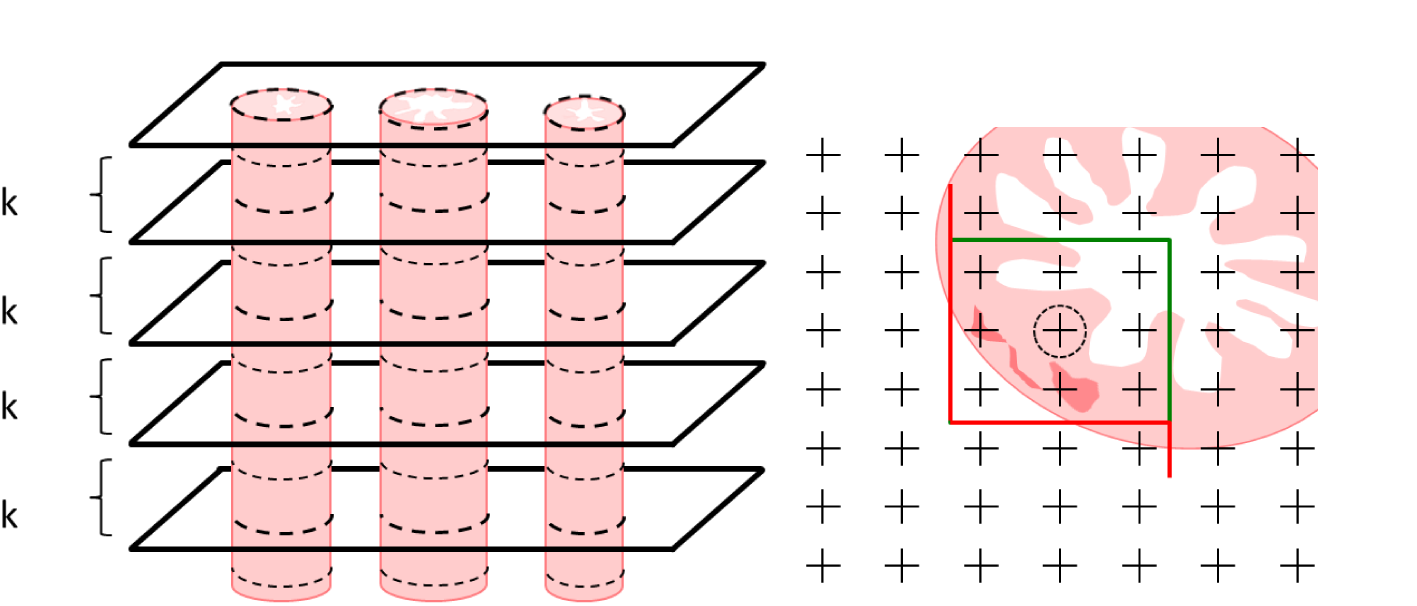
Schematic outline of sectioning, counting grid, and frame for stereology. Left) Tissue sections with a set interval (k) are taken for analysis eg. every 20^th^ section. The first section to be used is determined by a random number table. Right) example of the counting frame surrounding the 3×3 grid used to estimate the relative degree of inflammation. In the example, 2/9 points hit inflamed tissue. The central circled point (p) is used for volume estimation; and has a constant associated area a(p). For ilea, 36 sections/animal were collected and for livers, the entire organ was sectioned.

**Suppl. Figure S4:**
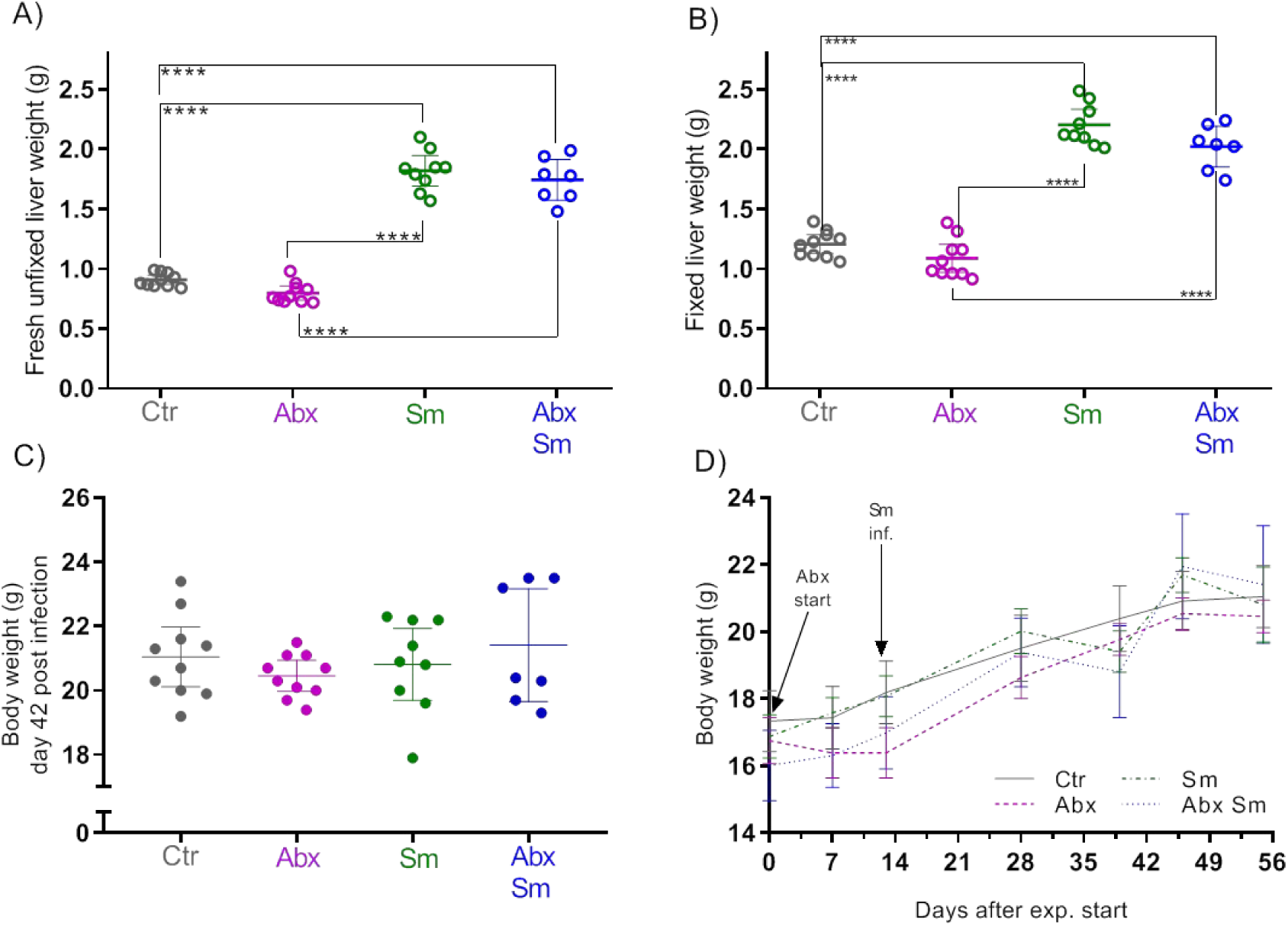
Liver and body weight. Weight (g) of A) fresh unfixed livers (analysis scales); B) 10% NBF fixed livers (tabletop scales); C) body weight of mice six weeks post infection and D) mean body weight of mouse groups over time (days) for control (Ctr), antibiotics treated (Abx), *S. mansoni* infected (Sm) and antibiotics treated and *S. mansoni* infected (Abx Sm) C57BL/6-NTac mice. Bars indicate means with 95%CI and stars the level of significance detected with ANOVA (Tukey’s multiple comparison’s test).

**Suppl. Figure S5:**
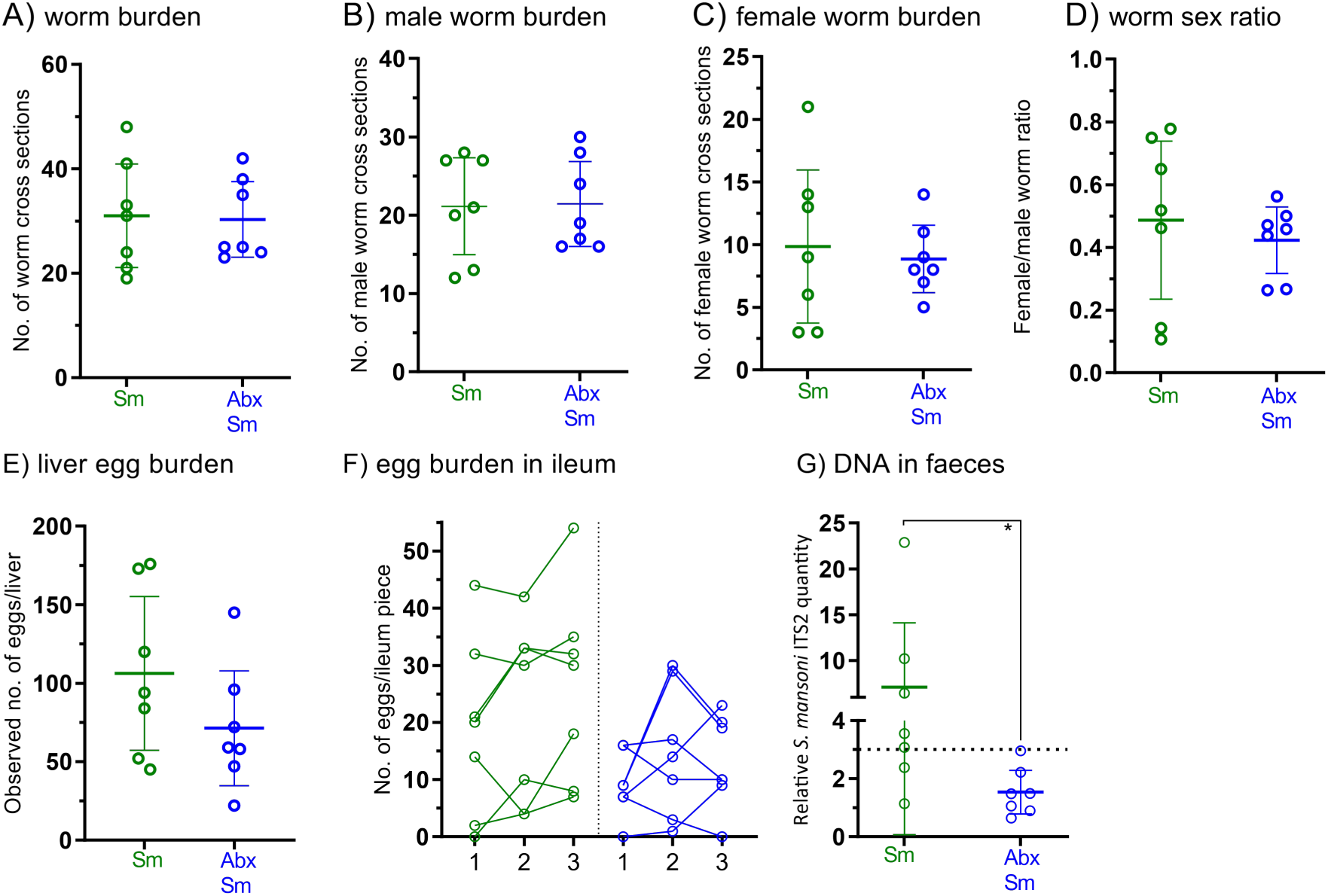
Infection intensity and egg dissemination. The worm burden is estimated by quantification of the number of worm cross sections for all slides sectioned for stereology per liver from *S. mansoni* infected groups (six weeks post infection; 120 cercs, n=7, 13-18 sections per liver) as shown in A) total worm sections, B) male worm sections, C) female worm sections and D) female/male worm section ratio. No differences in worm counts or macroscopic morphology of worms were observed from a small sample of perfused mice in a comparable model. E) show the total number of eggs present in all counting frames for each liver. F) The total number of eggs observed on 12 ileum cross sections for three ileum pieces per infected animal (36 sections/animal). Connecting lines depict counts for individual animals (1, 2, 3 where 1 is the most proximal, 2 is the middle and 3 is the most distal piece). No consistent pattern in the localization of eggs in terms of proximity to caecum or jejunum was observed. G) Illustrates the significantly lower relative *S. mansoni* DNA amount in faeces measured by ITS2 targeted RT-PCR (p=0.011, mean with 95%CI) in AbxSm mice compared to *S. mansoni* infected only (Sm).

**Suppl. Figure S6:**
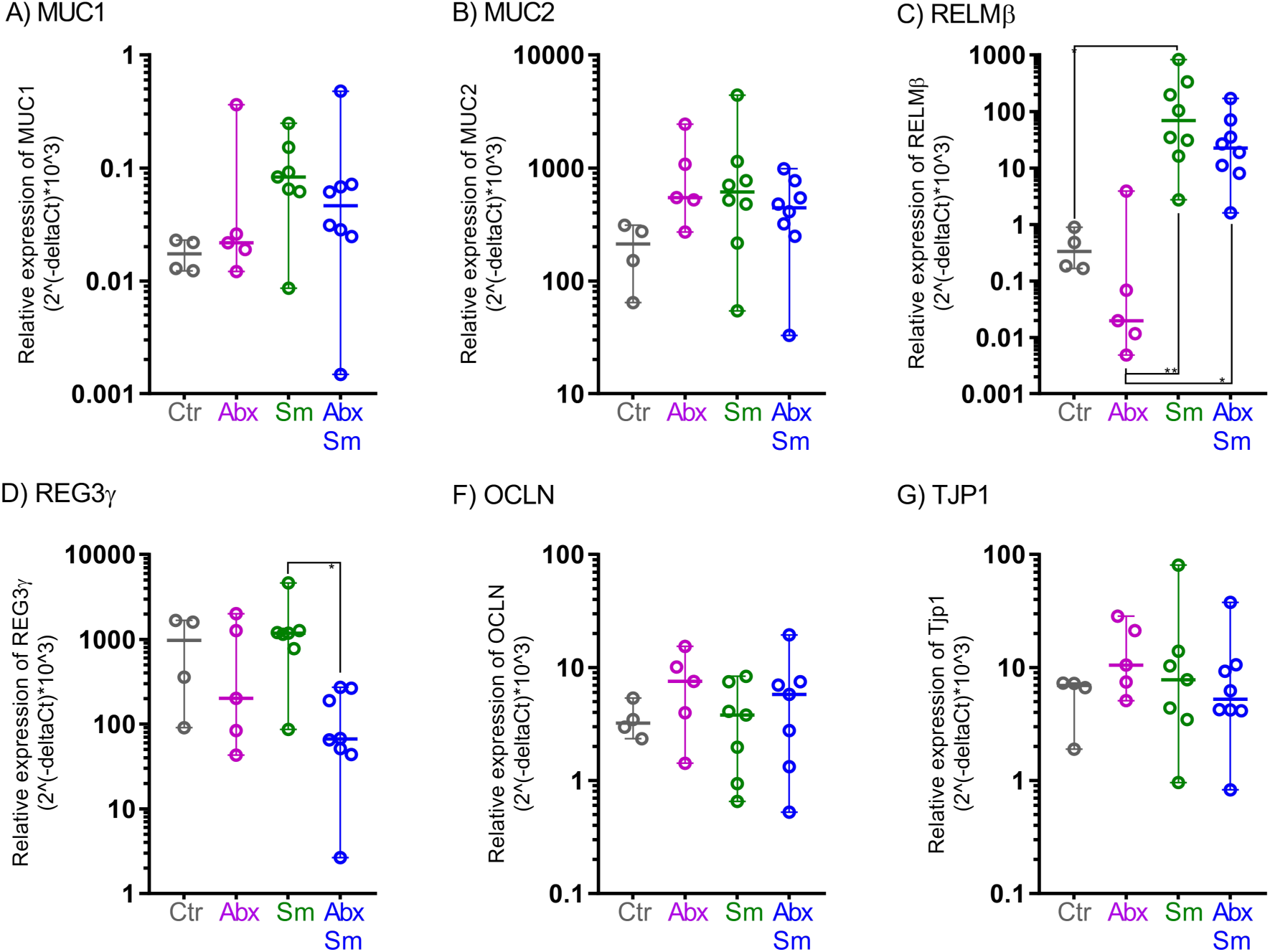
Relative gene expression of gut barrier function markers in ileum samples. Relative gene expression of the mucins A) MUC1 and B) MUC2; the antimicrobial peptide C) REG3γ; the cytokine D) RELM-β; and tight junction proteins E) OCLN and F) TJP1 encoding genes in ileum samples from control (Ctr), antibiotics treated (Abx), *S. mansoni* infected (Sm) and antibiotics treated and *S. mansoni* infected mice at eight weeks post infection. Each dot depicts the mean of duplicate measures in a single mouse. Asterisks indicate significant differences between groups (Kruskal-Wallis, Dunn’s multiple comparisons test).

**Suppl. Figure S7:**
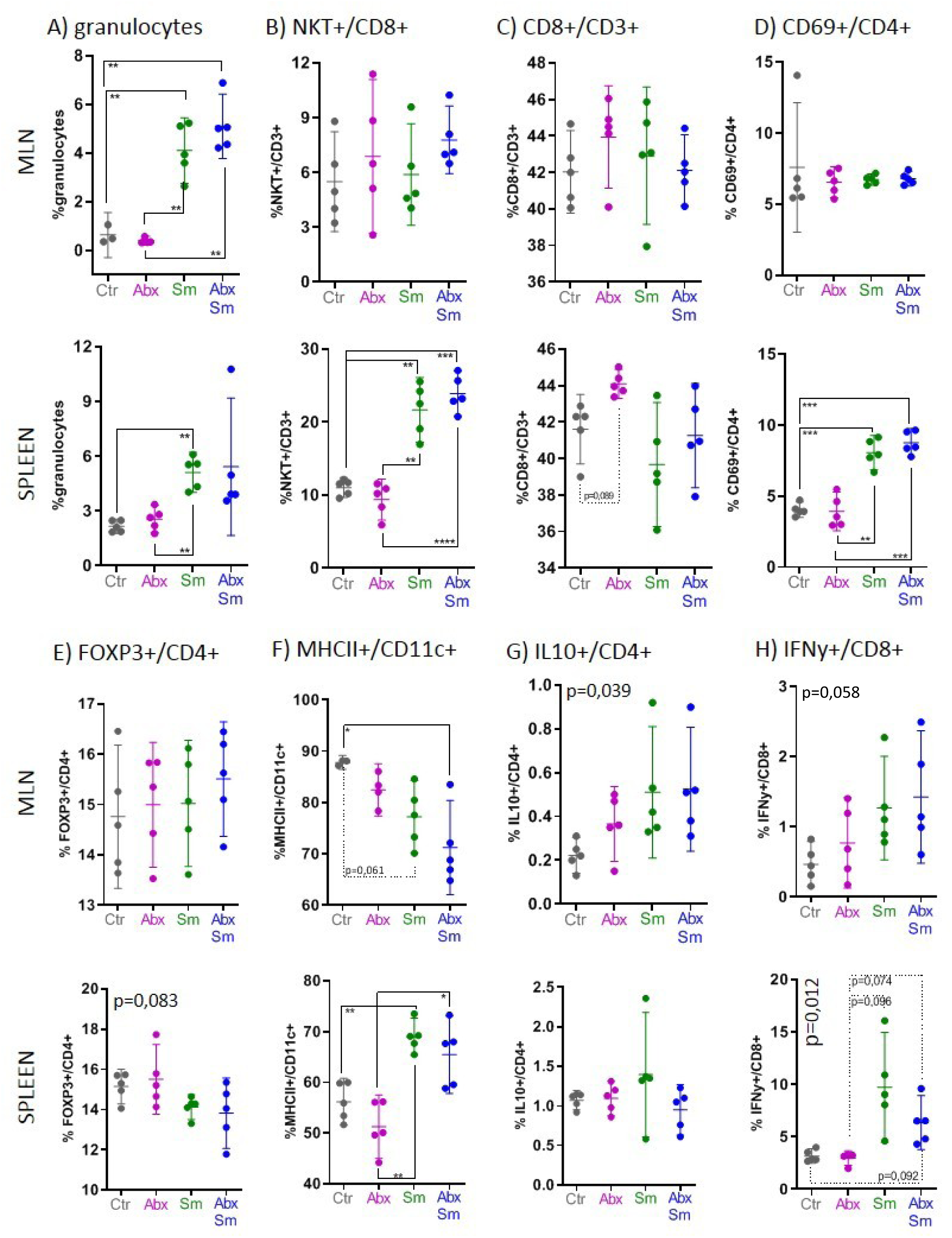
**Immune cell phenotypes in mesenteric lymph nodes and spleens by flow cytometry**. Graphs show proportions of cells positive for a range of markers as seen by flow cytometry. Cells were isolated from mesenteric lymph (MLN) and spleens from control (Ctr, n=5), antibiotics treated (Abx, n=5), *S. mansoni* infected (Sm, n=4), and antibiotics treated *S. mansoni* infected (AbxSm, n=5) mice six weeks post infection. MLN results are shown on top and spleen on the bottom for A) Granulocytes, B) NK-NKT+/CD3+, CD8+/CD3+, D) CD69+/CD4+, E) FOXP3+/CD4+, F) MHCII+/CD11c+, G) IL10+/CD4+ and H) IFNγ+/CD8+. Bars represent means with 95% CI. Stars indicate statistically significant differences between groups (Welch’s ANOVA with Dunnet’s correction for multiple testing, p<0,05 cut-off). All p values < 0,1 in group comparisons are indicated by dotted lines. A p value in the upper left corner means a significant difference detected by Welch’s ANOVA, but not by underlying pairwise testing.

**Suppl. Figure S8:**
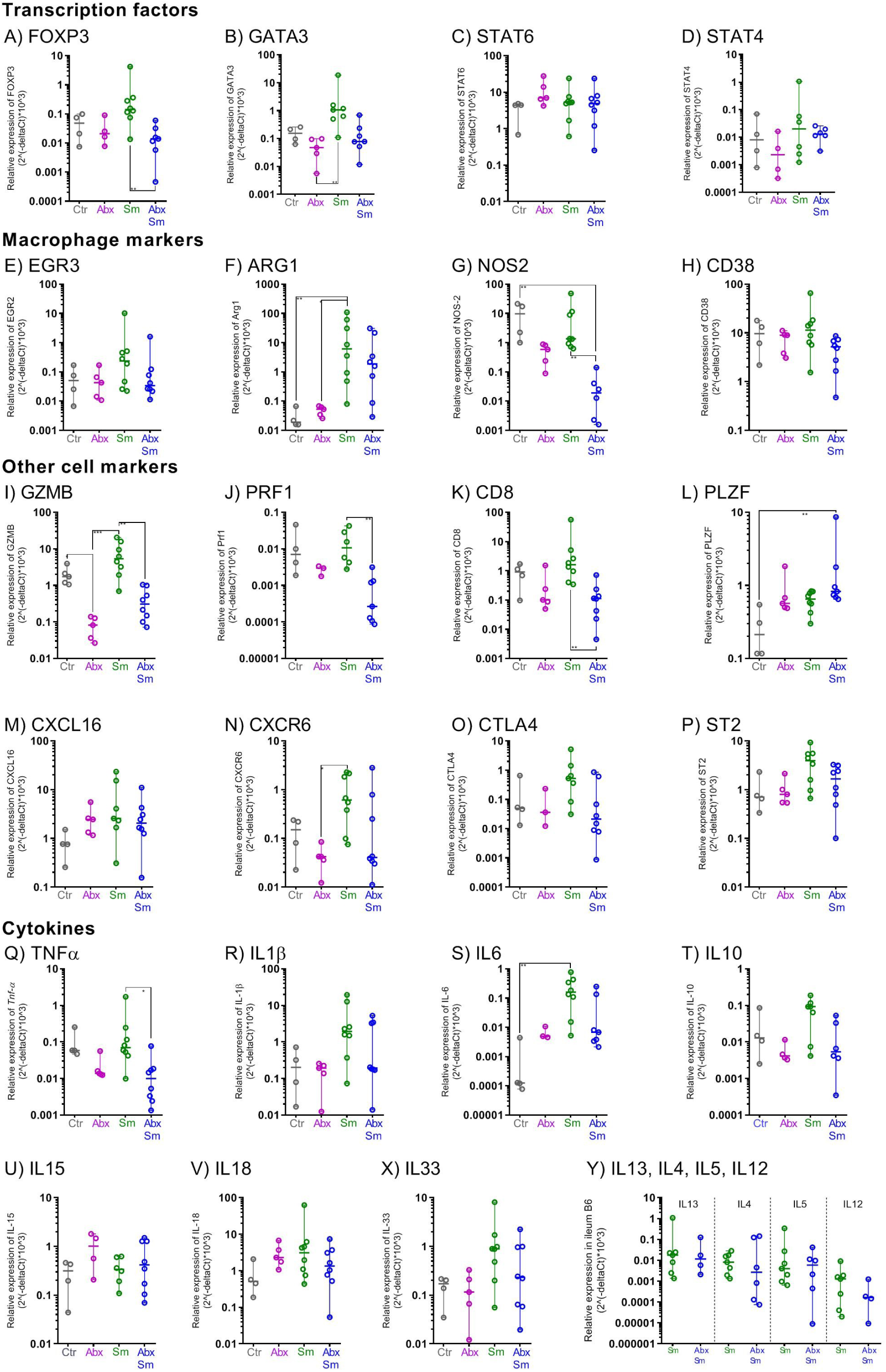
Relative transcription factor, cytokine, and cell marker expression in the ileum. Relative gene expression in ileum at eight weeks post infection of transcription factors A) GATA-3, B) FOXP3 C) STAT6, and D) STAT4; of macrophage phenotype markers E) EGR3, F) ARG1, G) NOS2 and H) CD38; of other markers I) GZMB, J) PRF1, K) CD8, L) PLZF, M) CXCL16, N) CXCR6, O) CTLA4 and P) ST2; of cytokines Q) TNFα; R) IL-1β, S) IL6, T) IL10, U) IL15, V) IL-18, X) IL33 and Y) IL13, IL4, IL5, IL12 for Control (Ctr), antibiotics treated (Abx), *S. mansoni* infected (Sm) and antibiotics treated *S. mansoni* infected (AbxSm) mice. Each dot depicts the mean of duplicate measures in a single mouse; the horizontal bar indicates the median with 95%CI. Asterisks indicate significant differences between groups (Kruskal-Wallis, Dunn’s multiple comparisons test). For Y) Ctr and Abx had too few mice with detectable signals.

**Suppl. Figure S9:**
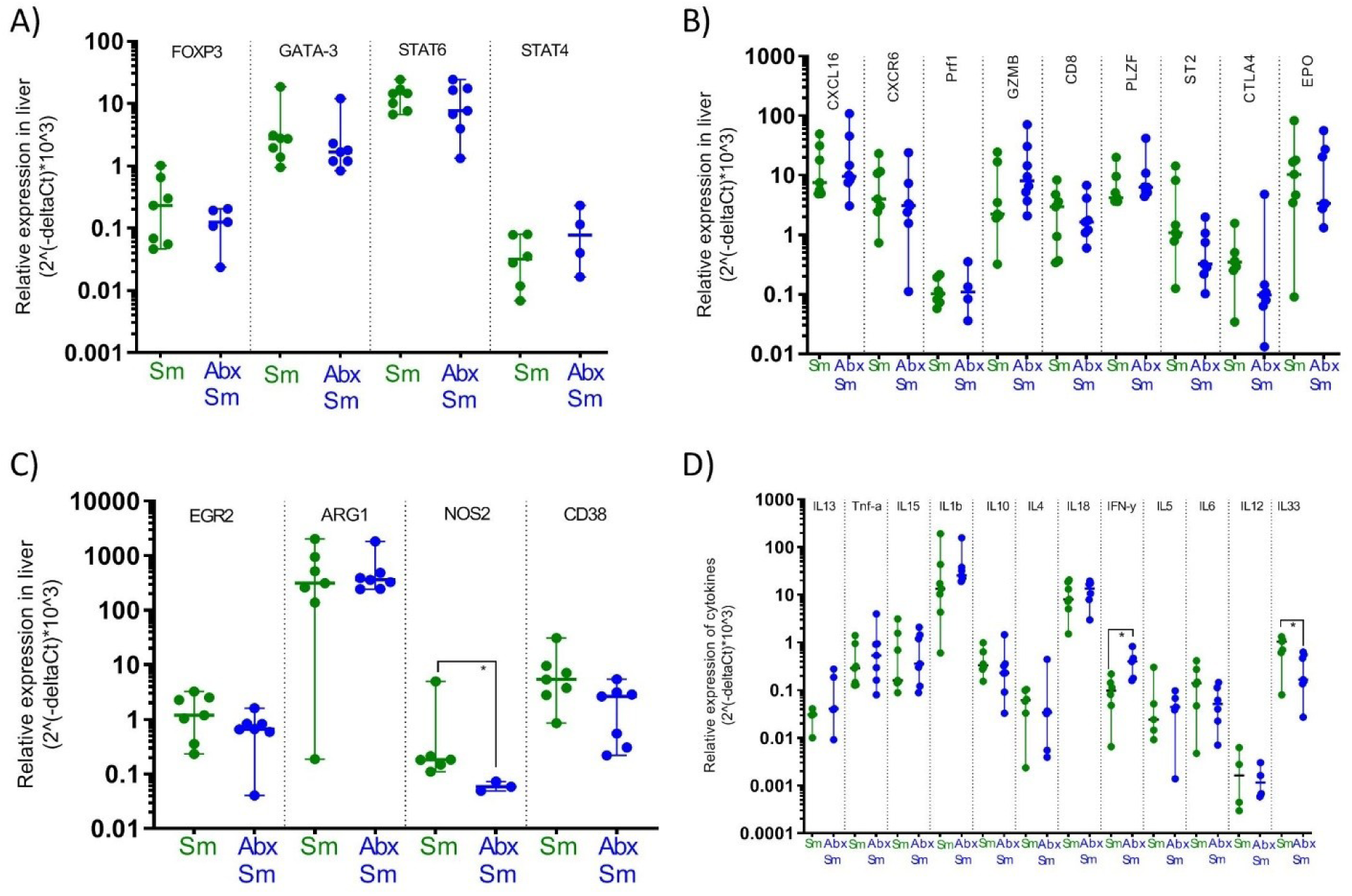
Relative transcription factor, cytokine, and cell marker gene expression in livers from *S. mansoni* infected mice. Relative gene expression in livers at eight weeks post infection of transcription factors FOXP3, GATA-3, STAT6 and STAT4; of other markers B) CXCL16, CXCR6, PRF1, GZMB, CD8, PLZF, ST2, CTLA4 and EPO; of macrophage phenotype markers EGR3, ARG1, NOS2 (significant difference even with outlier excluded) and CD38; and of cytokines IL13, TNFα, IL15, IL-1β, IL10, IL4, IL18, IFNy, IL5, IL6, IL12 and IL33 for Control (Ctr), antibiotics treated (Abx), *S. mansoni* infected (Sm) and antibiotics treated *S. mansoni* infected (AbxSm) mice. Each dot depicts the mean of duplicate measures in a single mouse; the horizontal bar indicates the median with 95%CI. Asterisks indicate significant differences between groups (Mann-Whitney).

**Suppl. Figure S10:**
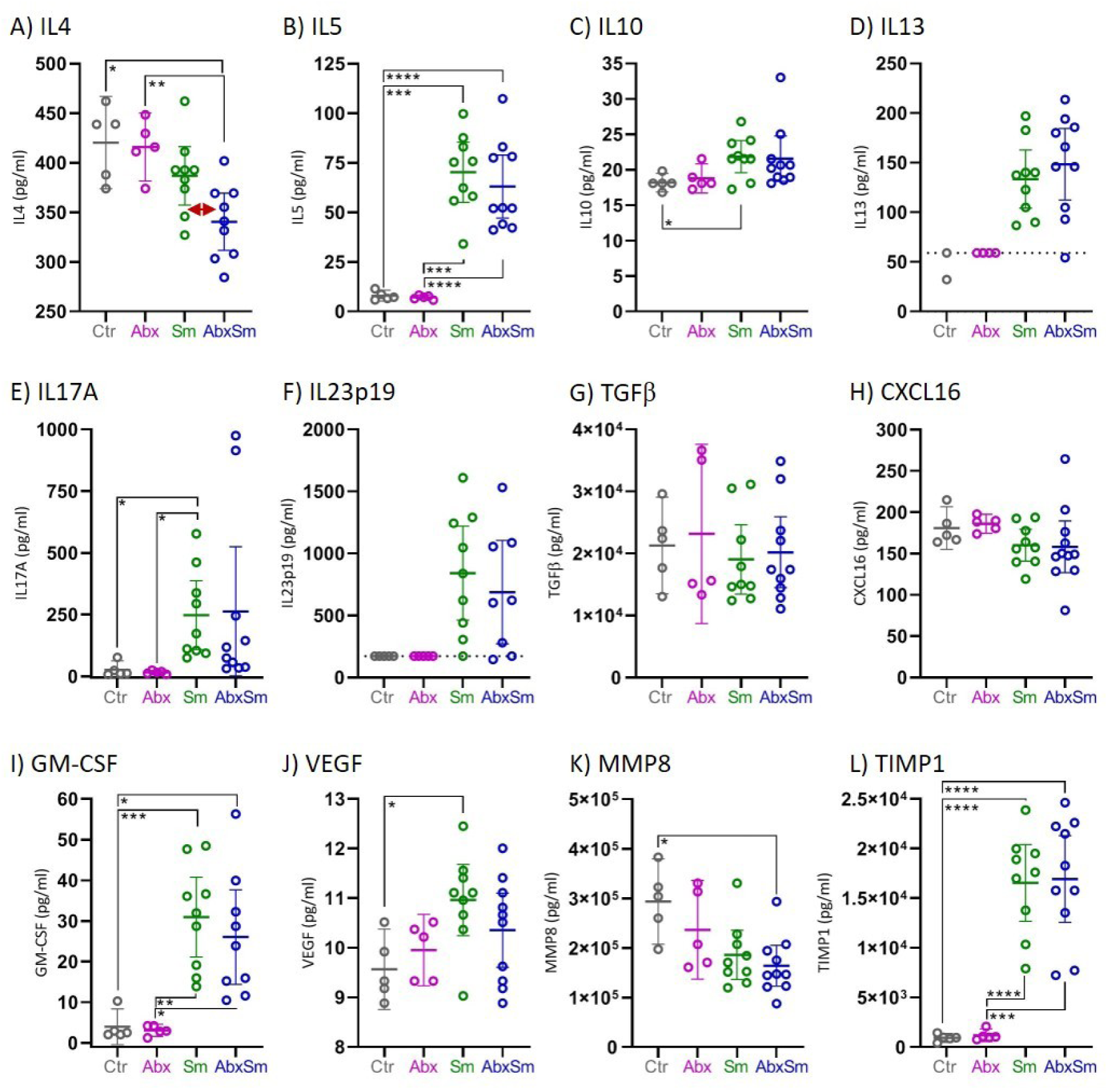
Immune parameters measured by Luminex. Levels (pg/ml) of cytokines A) IL4 (a single extreme outlier at 891 in AbxSm was excluded), B) IL5, C) IL10, D) IL13, E) IL17A, F) IL23p19 (a single extreme outlier at 4386 in AbxSm was excluded), G) TGFβ, H) CXCL16 and I) GM-CSF; growth factor J) VEGF; metalloproteinase K) MMP8 and TIMP metallopeptidase inhibitor 1 L) TIMP1 measured by Luminex magnetic bead assay for control (Ctr), antibiotics treated (Abx), *S. mansoni* infected (Sm) and antibiotics treated *S. mansoni* infected (AbxSm) mice eight weeks post infection. Bars are means with 95%CI. Stars indicate significant differences between groups by Welch’s ANOVA, Dunnett’s multiple comparisons test. For A) Sm and AbxSm groups are significantly different (p=0.020; T-test Welch’s correction, red horizontal arrow). Horizontal dotted line on D) and F) indicates that both Ctr and Abx groups had respectively IL13 and IL23 levels below the detection limit.

**Supplementary Table S1:**
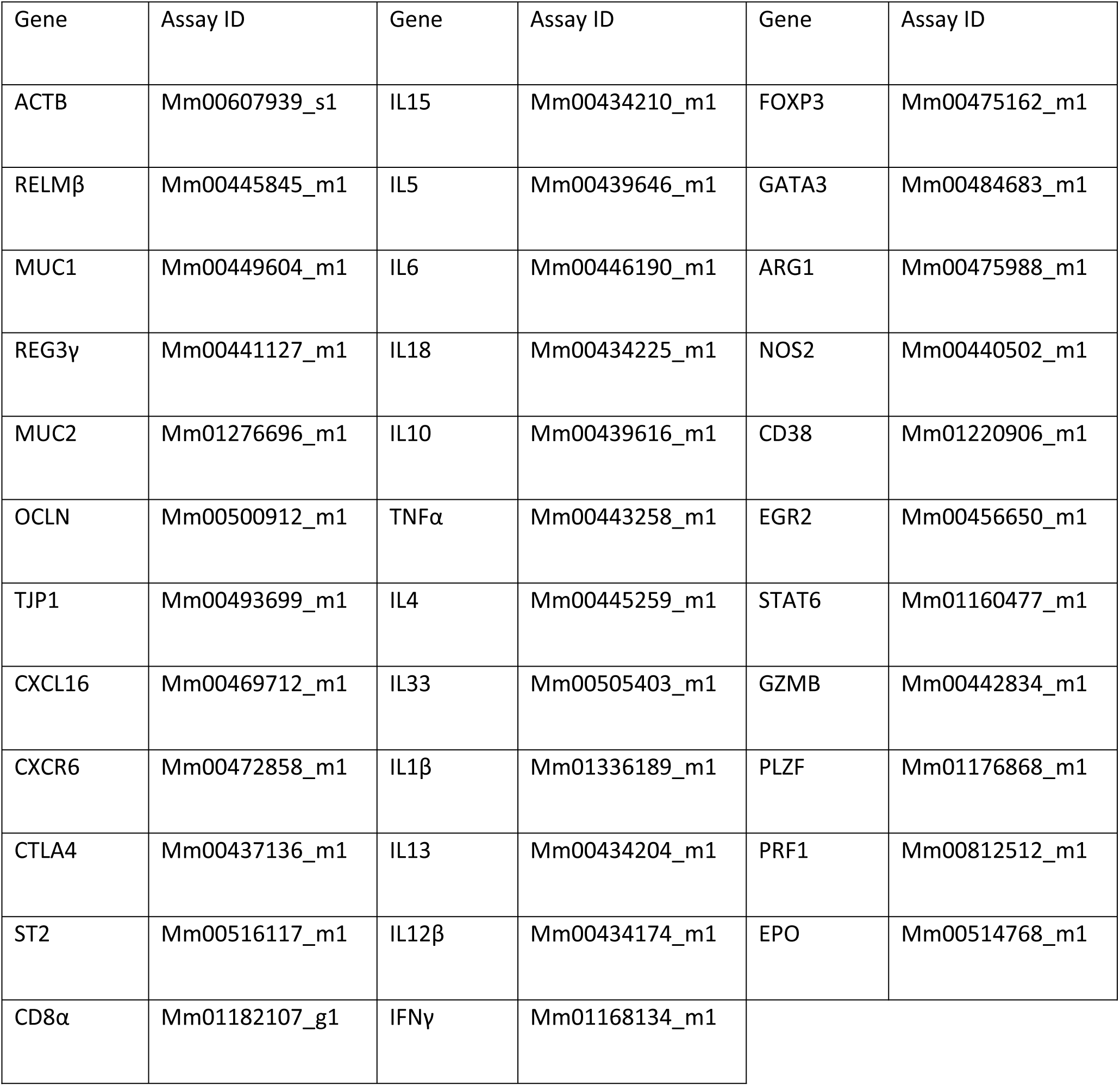
Taqman Gene Expression Assays (Applied Biosystems) used for Fluigdigm-based real-time PCR.

| Gene | Assay ID | Gene | Assay ID | Gene | Assay ID |
| --- | --- | --- | --- | --- | --- |
| ACTB | Mm00607939_s1 | IL15 | Mm00434210_m1 | FOXP3 | Mm00475162_m1 |
| RELMβ | Mm00445845_m1 | IL5 | Mm00439646_m1 | GATA3 | Mm00484683_m1 |
| MUC1 | Mm00449604_m1 | IL6 | Mm00446190_m1 | ARG1 | Mm00475988_m1 |
| REG3γ | Mm00441127_m1 | IL18 | Mm00434225_m1 | NOS2 | Mm00440502_m1 |
| MUC2 | Mm01276696_m1 | IL10 | Mm00439616_m1 | CD38 | Mm01220906_m1 |
| OCLN | Mm00500912_m1 | TNFα | Mm00443258_m1 | EGR2 | Mm00456650_m1 |
| TJP1 | Mm00493699_m1 | IL4 | Mm00445259_m1 | STAT6 | Mm01160477_m1 |
| CXCL16 | Mm00469712_m1 | IL33 | Mm00505403_m1 | GZMB | Mm00442834_m1 |
| CXCR6 | Mm00472858_m1 | IL1β | Mm01336189_m1 | PLZF | Mm01176868_m1 |
| CTLA4 | Mm00437136_m1 | IL13 | Mm00434204_m1 | PRF1 | Mm00812512_m1 |
| ST2 | Mm00516117_m1 | IL12β | Mm00434174_m1 | EPO | Mm00514768_m1 |
| CD8α | Mm01182107_g1 | IFNγ | Mm01168134_m1 | | |

**Suppl. Table S2:** LC-MS binary solvent elution gradient.

| Time [min] | Solvent A [%] | Solvent B [%] |
| --- | --- | --- |
| 0-2 | 85 | 15 |
| 2-11 | 85->45 | 15->55 |
| 11-12 | 0 | 100 |

